# Inversions maintain differences between migratory phenotypes of a songbird

**DOI:** 10.1101/2021.04.05.438456

**Authors:** Max Lundberg, Alexander Mackintosh, Anna Petri, Staffan Bensch

**Affiliations:** Department of Biology, Lund University, Sweden; Institute of Evolutionary Biology, University of Edinburgh, UK; Science for Life Laboratory, Uppsala Genome Center, Uppsala University, Sweden

## Abstract

Long-distance migration requires adaptations in a suite of behavioral, physiological and morphological traits. However, almost nothing is known about the genetic basis of these adaptations. The willow warbler *Phylloscopus trochilus* occurs in Europe with a northern and southern subspecies that show drastically different migration routes and wintering areas. Previous studies have demonstrated that the subspecies are genetically extremely similar except for three divergent chromosome regions, of which two are associated with the differences in migratory phenotypes and one is associated with an environmental gradient. Here we use a combination of long-read sequencing, linked-read sequencing and optical mapping to construct more complete and contiguous assemblies for both of the subspecies. We find evidence for inversions in each of the three divergent regions, which range from 0.4 to 13 Mb in size, and that breakpoints are associated with tandem repeat arrays or segmental duplications. The divergence times between inverted and non-inverted haplotypes are similar across the regions (∼1.2 Myrs), which is compatible with a scenario where the inversions arose in either of two allopatric populations that subsequently hybridized. The improved genome assemblies and annotation also allowed us to detect additional functional differences in the divergent regions that provide candidate genes for migration and local adaptations to environmental gradients.

## Introduction

For many birds, migratory behavior has a strong genetic component that is believed to be encoded as a set of innate migratory directions^1^ and a timing program^2^. Long-distance migration is also expected to lead to adaptations in morphological traits, such as wing shape^3^, and in physiological traits, such as optimally storing and using fat^4^. However, to date, very little is known about specific genes or pathways that underlie differences between migratory phenotypes^5^.

The willow warbler *Phylloscopus trochilus* is represented by two differentially migrating populations in Europe^6, 7^. The southern migratory phenotype (ssp. *trochilus*) occurs in Western Europe and migrates to Western Africa. The northern migratory phenotype (ssp. *acredula*) breeds in Northern Scandinavia and Eastern Europe and winters in Eastern or Southern Africa. The subspecies are otherwise morphologically and ecologically similar^8, 9^.

In the most comprehensive genetic study of the willow warbler to date, Lundberg et al.^10^ assembled a draft genome based on short-read data and used whole-genome resequencing and a customized 4000 SNP array to explore genetic differences between the migratory phenotypes. The vast majority of variants that were highly differentiated between the migratory phenotypes were located in three divergent regions on chromosomes 1, 3 and 5. Variation in the regions on chromosome 1 and 5 was strongly associated with migratory phenotypes while the region on chromosome 3 showed a stronger association with latitude and altitude. The clearly delimited plateaus of high genetic differentiation and the apparent lack of recombination between divergent southern and northern haplotypes in these regions suggested the presence of inversion polymorphisms. Loci underlying local adaptation and speciation have been found to be concentrated in inversions across many species of animals and plants^11^. By capturing co-adapted variants at linked loci, inversions facilitate the formation of supergenes, where complex phenotypes are inherited as if they were encoded by a single gene^12^. For example, inversion polymorphisms have been associated with different mating types in birds^13, 14^, social polymorphisms in insects^15^ and differences in migratory phenotypes in fish^16, 17^. In the willow warbler genome, however, no inversion breakpoints could be identified, presumably because they were located in repeat rich regions. In addition, the two migration-linked regions on chromosome 1 and 5, were split into two and ten scaffolds, respectively, making it difficult to know if the gene order within these regions is different from what is found in other birds.

Here we used long-read sequencing, linked-read sequencing, optical mapping and RNAseq to create a more complete, contiguous and well-annotated reference genome of a northern willow warbler. The new reference genome allowed us to explore the organization of the divergent chromosome regions, examine if additional highly differentiated regions between the migratory phenotypes might reside in parts of the genome not included in the previous short-read genome assembly, and to assess functional consequences of highly differentiated variants. To explore large-scale structural differences between the subspecies we also used linked reads and optical mapping to create a *de novo* assembly for a southern willow warbler. Finally, to elucidate the evolutionary history of each divergent region we fit models of population divergence using information within the blockwise site frequency spectrum (bSFS)^18^. This approach, adapted from Lohse et al.^19^, explicitly estimates the ancestral effective population size as well as the rate of migration, and so provides more accurate estimates of population divergence time than summary statistics where these parameters are assumed or ignored. If the divergent haplotypes in each of the regions, which are presently associated with each subspecies, are associated with inversions that arose independently within a single ancestral population, we would have no expectation of synchronized divergence times between regions (Figure 1). An alternative scenario is that the divergent haplotypes are a consequence of an ancient hybridization event, which has previously been hypothesized to explain the diversity in the extant willow warbler populations^20^. In this scenario, northern and southern haplotypes were unique to either of the hybridizing populations and were protected from recombining with each other through inversions, whereas the rest of the genome was homogenized through gene flow. In this case, we expect the divergence times to be similar across the regions (Figure 1), because they would not represent the actual inversion events, but rather the time of the ancient population split^21^.

**Figure 1.**
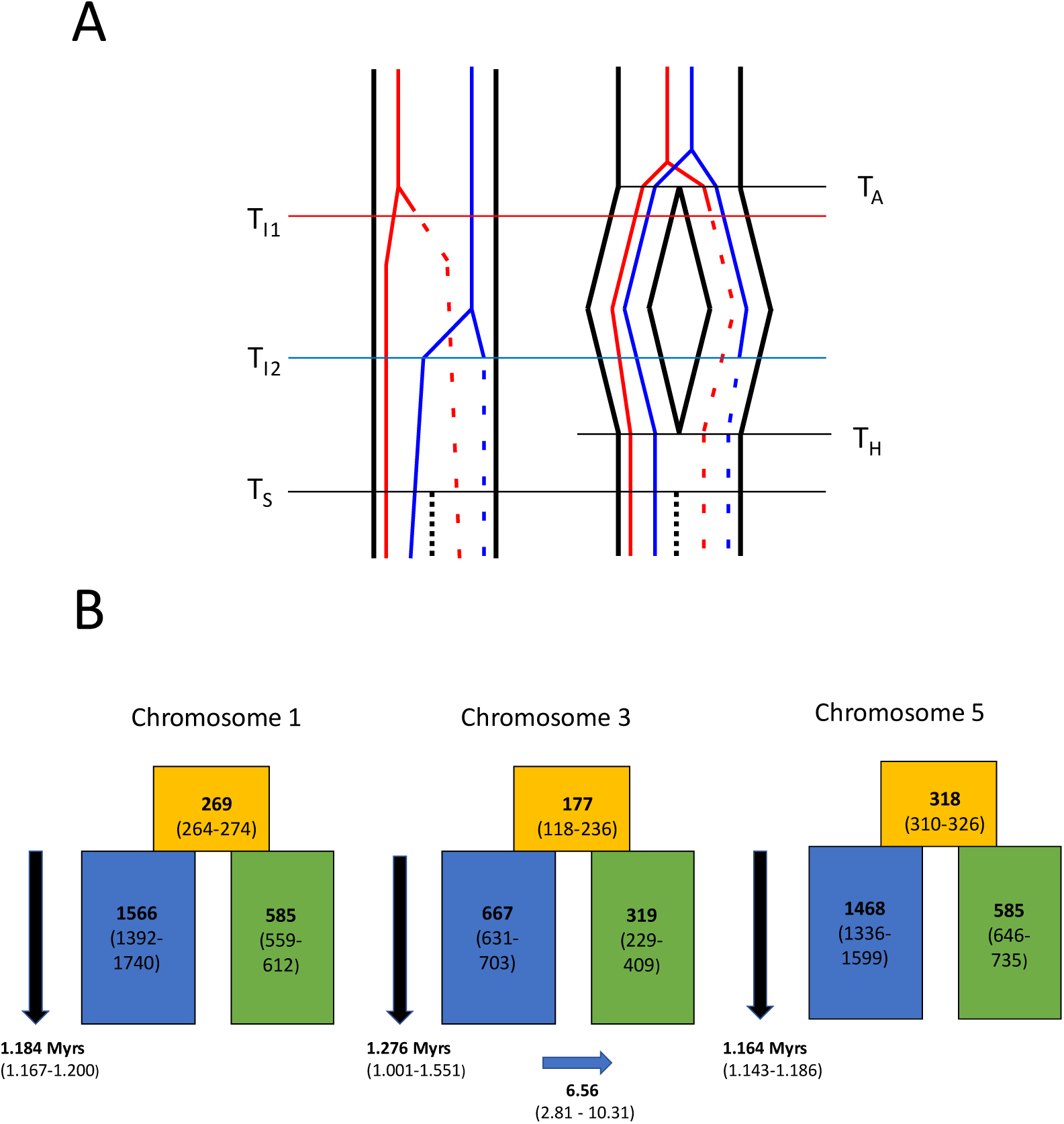
Evolutionary history of inversions. (A) Alternative population scenarios for two inversions (lines changing from solid to dotted) appearing at separate loci (red and blue) at timepoints T_I1_ and T_I2_, respectively. In the left scenario, the inversions appear as polymorphisms within a single ancestral population and eventually get sorted into one of two subspecies (T_S_). The divergence times of the inverted and non-inverted haplotypes will in this case reflect the timepoints of the inversion events. In the right scenario, an ancestral population splits into two allopatric populations (T_A_) and inversions appear in one of them. The two populations later come into secondary contact and merge (T_H_), before the inversions get sorted into one of two subspecies (T_S_). In this scenario, the divergence times of the inverted and non-inverted haplotypes for the two loci will be similar as they do not reflect the time of the inversion events, but the time of the ancestral population split (T_A_). (B) Population modelling parameter estimates (maximum composite likelihood with 95CI) for the three divergent regions in the willow warbler. Orange, blue and green boxes refer to ancestral, northern and southern effective population sizes, respectively, with numbers given in units of 10^3^. For chromosome 3, simulations confirmed a better fit for a model including migration (IM_2_), and in this case the blue arrow indicates the direction of migration with the rate given in units of 10^-7^. All three divergent regions have similar split times (∼1.2 Myrs), as expected from the right scenario in (A).

## Results

### Genome assemblies

Using a combination of long-read sequencing, linked-read sequencing and optical mapping we obtained a *de novo* assembly for a northern willow warbler consisting of 496 scaffolds with a N50 and L50 of 17.1 Mb and 17, respectively, and a contig N50 of 9.3 Mb (Supplementary Table 1). These contiguity metrics represent an order of magnitude improvement compared to those of a previous short-read willow warbler genome assembly^10^. The quality of the assembly was also verified by the presence of a high percentage of complete and a low percentage of duplicated single-copy bird orthologues (94.1 % and 1.1 % of 4,915 targeted genes, respectively, Supplementary Table 1).

The assembly of the southern willow warbler, which was constructed from a *de novo* assembly of linked reads that was further scaffolded by optical mapping data, was comprised of 39,596 scaffolds, had a scaffold N50 and L50 of 12.4 Mb and 24, respectively, and a contig N50 of 129 kb (Supplementary Table 1).

The northern and the southern assembly contained 14.8 % and 12.9 % annotated repeats, respectively, compared to 9.5 % in the previous willow warbler assembly (Supplementary Table 2). The northern assembly was annotated with 18,915 protein-coding genes based on a combination of willow warbler RNAseq data (Supplementary Table 3) and protein data from other bird species.

### Differences between subspecies

We explored genetic differentiation between the subspecies by mapping whole-genome resequencing data of 11 southern and 11 northern willow warblers (Supplementary Table 4) to the northern reference genome. The weighted average F_ST_ between the northern and southern samples for 44 million bi-alllelic SNPs was 0.006. The number of highly-differentiated variants was extremely small, with almost all located in the previously identified divergent chromosome regions (Supplementary Table 5). For example, of 11,623 variants with F_ST_ ≥ 0.7, only 208 were found outside of the three regions. The majority (80 %) of these variants were found on 24 scaffolds (median size: 1,071,152 bp, range: 19,698 bp - 12,491,261 bp) that could not be assigned to specific chromosomes in the other bird species. These scaffolds contained a high proportion of repeats (85-93 % of the ungapped length), a high GC content (49-53 %) and mostly multi-copy genes (e.g. olfactory genes), although it is unclear to what extent these genes are functional. Additionally, coverage in the southern samples was generally lower than in the northern samples (Supplementary Figure 1). This could suggest that these scaffolds represent genomic regions that are unique to the northern subspecies in terms of repeat composition and that the mapped reads from the southern samples are either from homologous but differently organized repeat-rich regions or from similar repeats in other parts of the genome.

### Chromosome 1 region

In the northern assembly, the highly differentiated region on chromosome 1 was assembled into an 11.7 Mb scaffold that consists of two contigs separated by a gap of 58 kb (Figure 2). The scaffolding of the contigs was supported by 10x chromium data and is concordant with the orientation found in the collared flycatcher and zebra finch genome assemblies. The gap is surrounded by arrays of a 413 bp tandem repeat (9.6 kb on the left and 0.5 kb on the right side of the gap, respectively; Supplementary Figure 2) that are lacking in the corresponding intervals in the genomes of the zebra finch and the collared flycatcher. This tandem repeat is classified by repeatmodeler as a part of a 2,021 bp DNA transposon CMC-EnSpm repeat, which is annotated in 313 intervals on 87 scaffolds. More similar and complete copies of the tandem repeat (≥90 % identity and ≥50 % of length), however, are restricted to the divergent regions of chromosomes 1 and 3 and on the scaffolds surrounding them based on synteny to other bird genomes.

**Figure 2.**
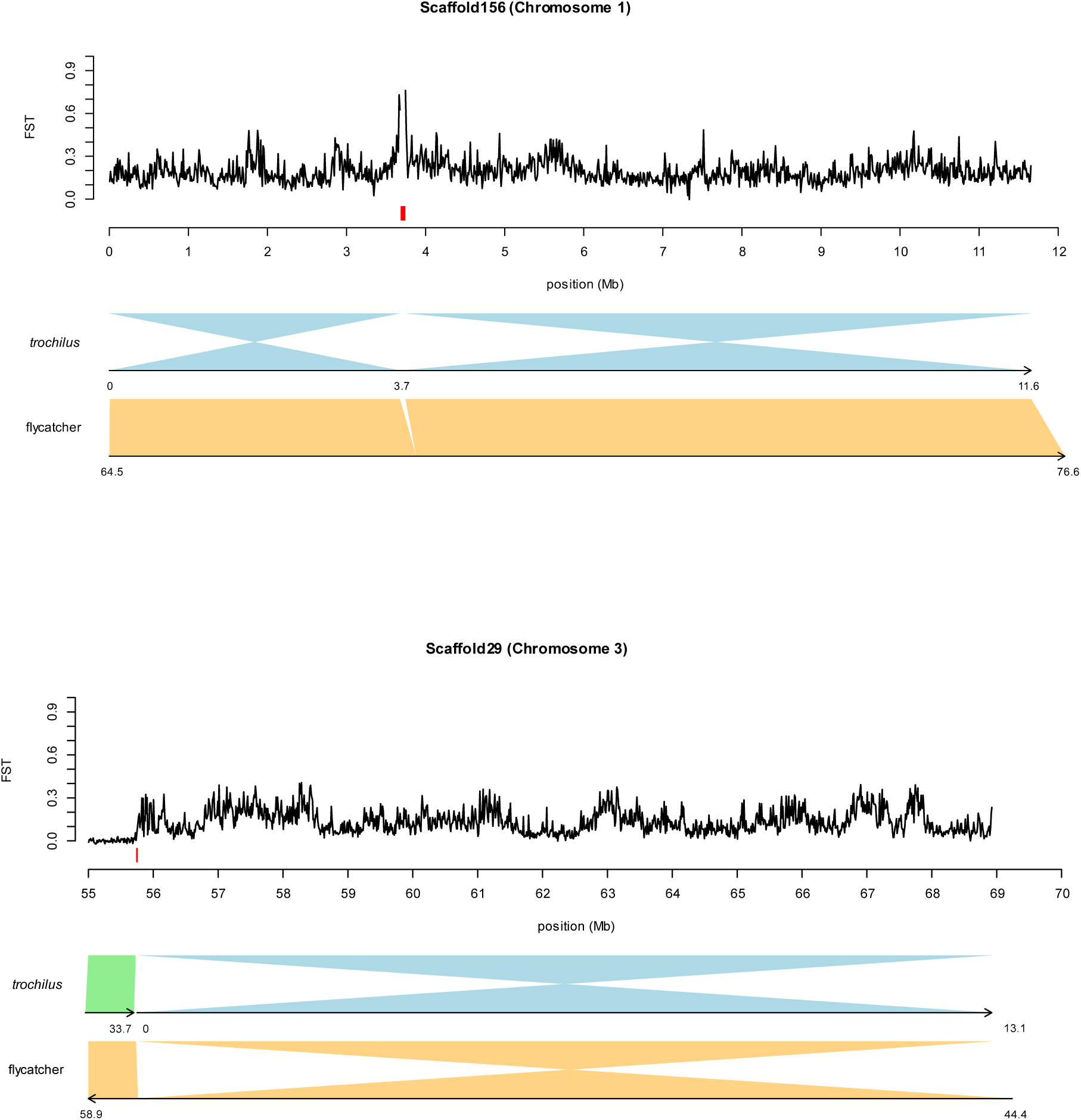
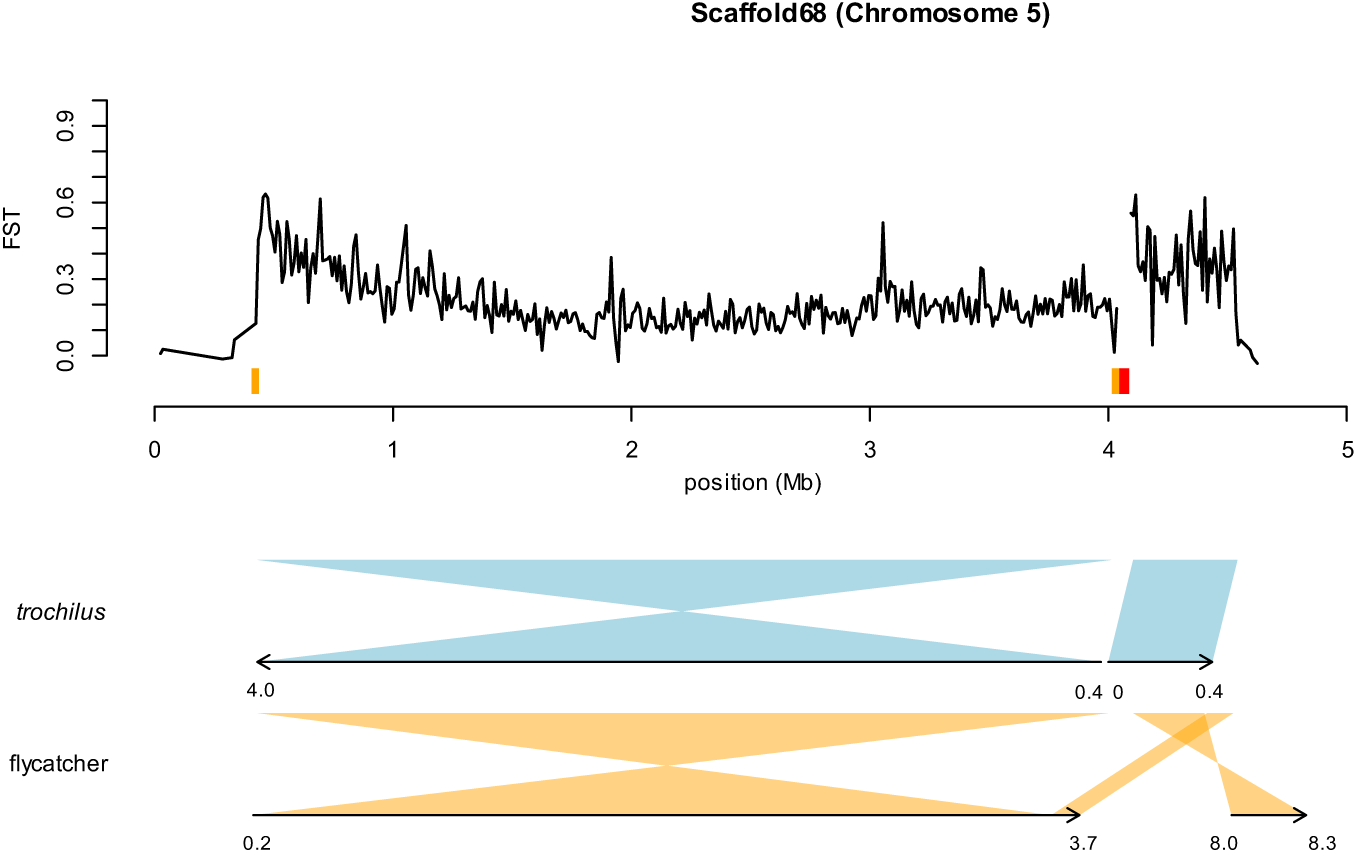
Divergent regions on chromosome 1, 3 and 5 in the northern assembly. The top panel shows genetic differentiation (weighted F_ST_) between 11 resequenced samples from each subspecies for bi-allelic SNPs in 10 kb non-overlapping windows. Red segments show the location of large gaps, which in the region on chromosome 1 and 3 are associated with tandem repeats. Orange segments show the location of 31 kb duplicated intervals in the chromosome 5 region. Lower panels show synteny between the northern reference genome and the southern genome (light blue and green) and the collared flycatcher genome (orange), respectively. For each comparison, the bottom line represents the position (Mb) in the other genome. The zebra finch genome shows the same major structural differences as seen in the flycatcher genome and has therefore not been included.

The differentiated region could not be joined with other scaffolds on chromosome 1. In both the collared flycatcher and the zebra finch, the orientation of the surrounding region is conserved and the ends of the divergent scaffold are expected to be located around 1 and 10 kb, respectively, from the ends of the adjacent scaffolds. The start of the divergent scaffold has a 1.9 kb array of the same tandem repeat as present in the large gap (Supplementary Figure 2) and is missing in the other species. This suggests that the difficulty to further scaffold the divergent region in the willow warbler may be associated with potentially large arrays of tandem repeats.

The southern *de novo* assembly also contained a single scaffold (11.6 Mb) for this region. Compared to the genome of the northern willow warbler and other bird genomes, the southern genome shows two tandem inversions (Figure 2). These two inversions, which are located on either side of the gap in the northern assembly, are 3.7 Mb and 7.9 Mb in length. The difference in breakpoints between the southern and the northern sample was also supported by alignments of their linked reads and optical maps (Supplementary results, Supplementary Figure 3).

Genetic differentiation between 11 resequenced samples from each subspecies was high across the entire region (mean weighted F_ST_ in 10 kb windows: 0.19), but showed prominent peaks overlapping the breakpoint region in the central part of the scaffold (Figure 2).

### Chromosome 3 region

In the northern reference assembly, the highly differentiated region on chromosome 3 was assembled into a single scaffold of 13.2 Mb that was further scaffolded (based on linked read data) with a 55.7 Mb undifferentiated part of chromosome 3, but in the opposite direction as predicted from the collared flycatcher and the zebra finch genomes (Figure 2). Hence, this suggests that the northern willow warbler has a derived inversion in the region. The gap separating the differentiated and undifferentiated regions is estimated to be 21 kb and is surrounded by 4.1 and 8.1 kb of the same type of tandem repeat as identified in the chromosome 1 region. As also seen on chromosome 1, there is no indication that this tandem repeat array is present in the same interval in the genomes of zebra finch and collared flycatcher. Based on the other two bird genomes, the other end of the differentiated and inverted region is expected to be 1 kb from its adjacent scaffold. The start of the adjacent scaffold contains 3 kb of the same tandem repeat as seen at the other end of the differentiated region (Supplementary Figure 2).

In the southern *de novo* assembly, the differentiated region was also assembled into a single scaffold of 13.2 Mb. The scaffold could be aligned over its full length to the northern reference genome, but did not show any structural differences suggestive of a rearrangement.

Highly differentiated variants were present across the entire differentiated region (mean weighted F_ST_ in 10 kb windows: 0.14), but as opposed to the region on chromosome 1, we did not see any clear breakpoint effect (Figure 2).

### Chromosome 5 region

In the northern reference assembly, the highly differentiated region on chromosome 5 was assembled into a single scaffold of 4.6 Mb (Figure 2). The differentiated part of this region (4.1 Mb) is surrounded by 430 and 80 kb of repeat-rich regions that could not be assigned to any chromosome in the zebra finch or collared flycatcher genome.

The southern assembly contained a 4 Mb scaffold covering the differentiated region but not any undifferentiated parts of the genome (Figure 2). Relative to the southern scaffold, the northern scaffold shows a 3.6 Mb inverted interval and a 445 kb collinear interval that are separated by a 41 kb gap (Figure 2). The collared flycatcher and zebra finch also shared the inverted interval with the southern willow warbler (Figure 2), although the breakpoints are not identical, which suggests that the northern subspecies has a derived rearrangement. Based on the mix of inverted and collinear intervals, a possible scenario is that the entire differentiated interval was first inverted in the northern subspecies and a second inversion event restored the order at the end.

Alignments of linked reads and the optical maps also suggested the presence of breakpoints at the start of the divergent region (∼ 400 kb) and immediately downstream of the 41 kb gap (∼4 Mb) (Supplementary Results; Supplementary Figures 3, 4). Within the breakpoint region at the start, there is a 31 kb segmental duplication that shows a 94 % similarity to an interval (in the same orientation) upstream of the breakpoint associated with transition between the inverted and collinear interval (Figure 2). The duplicated interval is rich in repeats and contains a truncated copy of the Nucleolar pre-ribosomal-associated protein 1 (URB1) gene, which in birds is located on chromosome 1. The copies associated with the duplicated interval contain a few exons found at the beginning and towards the end of the complete gene.

Genetic differentiation between resequenced southern and northern birds was high (mean weighted F_ST_ in 10 kb windows: 0.21) within the entire region, but was on average higher in the second part of the region associated with the interval that is collinear between subspecies (Figure 2). As in the case of the region on chromosome 1, there was a breakpoint effect in genetic differentiation, with particularly high values close to the start of the differentiated region (∼0.4 Mb) and the start of the inverted interval (∼4.1 Mb).

### Functional differences in the divergent regions

With the improved assembly and annotation we examined potential functional differences between the subspecies in the three chromosome regions. None of the breakpoint intervals overlapped with or were very close to an annotated functional protein-coding gene (range: 1.6-67.3 kb, Supplementary Table 6). We also explored whether any SNPs or short indels with high differentiation (F_ST_≥0.7) between southern and northern homozygotes in each of the divergent region were predicted to have a moderate to high effect on protein-coding genes. Across the three regions we found 66 nonsynonymous mutations and an in-frame insertion located in 45 genes (Supplementary Table 7). We additionally found one frameshift mutation in general transcription factor IIIA (GTF3A), which is located within the region on chromosome 1. This change, which represents a derived deletion in the northern subspecies, modifies four amino acids at the end and further extends the protein with three amino acids. Although the genes with protein-coding changes were functionally diverse, some shared more specific functions. In particular, nonsynonymous mutations were found in three fatty acid desaturase genes (FADS2, FADS1L1, FADS1L2) that are located in tandem in the second differentiated interval on chromosome 5. Two of these genes (FADS2, FADS1L2) are also annotated as involved in “oxidation-reduction process” together with cytochrome b5 reductase 2 (CYB5R2) and gamma-butyrobetaine hydroxylase 1 (BBOX1), also on chromosome 5, and crystallin lambda 1 (CRYL1) on chromosome 1.

Finally, we searched for highly differentiated structural variants between southern and northern samples in the divergent regions. We detected 105 deletions (mean size: 103 bp, range: 17-1,495 bp) and 68 insertions (mean size: 31 bp, range: 15-341 bp) that had a F_ST_ ≥ 0.7 between southern and northern homozygotes in the divergent regions. The majority of the structural variants (102/173) were located outside of the annotated genes with a median distance of 38 kb and most that were overlapping genes were found in relatively large introns (median 20 kb). Only two of the variants overlapped exons of protein-coding genes: a 21 bp insertion in the 3’ UTR in Interferon gamma receptor 1 (IFNGR1) located in the region on chromosome 3 and a 203 bp deletion in the 3’ UTR of the Stomatin Like 3 (STOML3) gene located in the region on chromosome 1.

### The age of divergent regions

We used two high-coverage resequenced samples of each subspecies to estimate the population divergence time between the subspecies in each of the divergent regions (Supplementary Table 4). First, we calculated net divergence (d_a_)^22^ and found similar values across the three regions (Supplementary Table 8). Using a germline mutation rate estimated from the collared flycatcher^23^, and assuming a generation time of 1.7 years^8^, these values correspond to divergence times of 487, 529, and 523 kyr for the regions on chromosomes 1, 3, and 5, respectively. As a complementary, but not independent measurement of divergence, we calculated relative node depth (RND)^24^ using a dusky warbler *Phylloscopus fuscatus* (Supplementary Table 4) as an outgroup. The RND estimates were similar across the regions (0.35-0.40, Supplementary Table 8) and suggest that divergence arose far more recently than the shared ancestor of the willow warbler and dusky warbler.

Divergence-based methods are expected to lead to underestimated divergence times if there has been migration occurring between populations. Furthermore, if diversity in the ancestral population was much larger or smaller than in the contemporary populations, d_a_ would be biased upwards or downwards, respectively. To overcome these caveats, and obtain more robust estimates of population divergence, we fitted demographic models involving divergence and gene flow between two populations for each divergent region. We used the software gIMble^25^, which leverages information within the blockwise site frequency spectrum (bSFS)^18^ to compare the support (composite likelihoods (CLs)) for different demographic models and parameter estimates.

For the divergent regions on chromosomes 1 and 5 the best fitting model (i.e. the highest CL) was the IM_1_ model. In this model the population split is followed by a constant rate of migration (m) from the southern population to the northern population forwards in time. For the divergent region on chromosome 3 the greatest CL was found using the IM_2_ model, where migration occurs in the opposite direction to IM_1_. However, the IM models are expected to always achieve a greater CL than strict isolation (SI) models because they include an additional parameter, the rate of migration. Additionally, the maximum composite likelihood (MCL) estimates of migration rate were small (4.46×10^-7^, 6.56×10^-7^ and 3.66×10^-7^ for chromosomes 1, 3, and 5 respectively), which, although consistent with limited recombination within inversions, suggests that the IM model may not fit significantly better than SI. Therefore, to test whether the IM models gave a significant improvement in CL, we simulated replicates under the optimized SI parameters for each divergent region and recorded the difference in CL between the IM and SI model. For the regions on chromosomes 1 and 5 we found that the improvement in CL between models was entirely consistent with a history of strict isolation (Supplementary Figure 5). By contrast, the improvement in CL observed for the chromosome 3 region is greater than we would expect if there had been no migration. Hereafter we present and discuss parameter estimates from the simpler SI model for the chromosome 1 and 5 regions and from the IM_2_ model for the chromosome 3 region (Figure 1).

The MCL parameter estimates suggest that the divergence time within each divergent region is around 1.2 Myr and thus far greater than was estimated by our d_a_ calculation. Effective population sizes were consistently estimated to be higher in the northern willow warblers and those of the ancestral populations smaller than in the contemporary populations. The similar parameter estimates among the three regions, especially divergence time (1.12-1.28 Myrs), suggests that they have a shared demographic history and supports a scenario where inversions happened in allopatric populations.

Consistent with the idea that almost all of the genetic differentiation between these subspecies is confined to only three regions, the MCL estimates of population divergence time presented above are an order of magnitude greater than analogous estimates over the rest of the genome (Supplementary Table 9). However, the distribution of coalescence times outside of the divergent regions may still contain information about the demographic history of these subspecies. For example, a sustained period of population structure will result in a reduced coalescence rate (increased N_e_) until populations become admixed again. To test whether the coalescence rate over time follows such a pattern, we used the sequentially Markovian coalescent (SMC) implemented in MSMC2^26^. For each sample, Ne was estimated to have been highest during the Pleistocene, peaking at ∼450 kya, and lower in more recent (<200kya) and distant (>2Mya) past. Although there are multiple explanations for this pattern, it is at least consistent with the possibility that a period of allopatry, beginning 1.2Mya and ending <450kya, is the reason that different divergent regions have a shared demographic history.

## Discussion

Using highly contiguous genomes we have demonstrated that the divergent regions separating the two willow warbler subspecies are associated with structural rearrangements. We also corroborated the results of Lundberg et al.^10^ by finding that virtually all the highly differentiated SNPs and indels between the subspecies are located in these regions. However, the long-read sequencing and optical mapping data enabled us to identify additional differences in previously overlooked repeat-rich scaffolds that may represent more large-scale structural differences between the subspecies. Due to their high repeat content, we failed to assign these scaffolds to specific chromosomes in other bird species, nor determine whether they are linked to any of the three previously identified divergent chromosome regions.

### Age and demography of the divergent regions

By fitting demographic models to the blockwise site frequency spectrum, we found that the northern and southern haplotypes have divergence times of ∼1.2 Myrs across the three regions (Figure 1). These estimates are considerably lower than the divergence time between the willow warbler and its closest relative, the chiffchaff, which is estimated to be around 5 Myrs^27^. Hence, we can reject the hypothesis that the presence of the divergent haplotypes within the willow warbler is a result of introgression from an extant *Phylloscopus* species.

Our analyses provide support for a previous hypothesis that the extant willow warbler is a result of an ancient hybridization event between two divergent subspecies or species^20^. The similar divergence times across the regions are compatible with a scenario where an ancestral population was split into two allopatric populations that subsequently hybridized with each other and homogenized the genome except for the divergent regions (Figure 1). In this scenario, the estimated divergence times would represent the time around the population split, and the structural rearrangements would have appeared at some time between the population split and the secondary contact event. The inverted haplotypes could have segregated at low frequency in either of the allopatric populations and increased in frequency at the time of secondary contact due to positive selection^21^. In this case, the structural rearrangements would have been selected for because they protected favorable allele combinations in the chromosome regions, for example those associated with adaptations to specific migratory routes in each population, from being broken apart due to gene flow and recombination^28^.

The genome-wide changes in effective population size over time as determined from the MSMC2 analysis (Supplementary Figure 6) are largely compatible with the proposed scenario of allopatric populations. The genome-wide effective population size could be increased when there is population structure^29^ and in willow warblers we observe an increase around the estimated divergence time of the northern and southern haplotypes. The decline in population size starting around 400 kya may then reflect the merging of the allopatric populations. However, we admit that the changes in genome-wide Ne could as well be caused by census population changes, although the scenarios are not mutually exclusive.

### Structural rearrangements in the divergent regions

In the divergent regions on chromosome 1 and 5, which are associated with differences in migratory phenotypes, we found evidence for rearrangements that are adjacent to or nested within each other (Figure 2). Complex rearrangements have been observed in a wide range of taxa^17, 30, 31^ and are likely to reduce gene flow even further between the divergent haplotypes. Alignments to other bird genomes enabled us to determine which of the subspecies has the derived or ancestral gene order within each region (Figure 2). For the region on chromosome 1, the southern subspecies has the derived gene order, whereas in the other two regions the northern willow warbler appears to possess derived rearrangements. The spread of the inverted haplotypes in either population at the secondary contact event could be expected to have given rise to a selective sweep that reduced variation^32^. However, we did not observe any consistent effect of inverted haplotypes on diversity, which suggests that any sweeps happened sufficiently long ago for diversity to accumulate and/or that the inverted haplotypes had been segregating some time before the selection event took place and generated softer sweeps. Instead, the northern haplotypes consistently showed higher diversity than the southern haplotypes and were assigned higher effective population sizes in the modelling analysis (Figure 2). The northern haplotypes are currently found over a larger geographical range than the southern haplotypes, particularly for the chromosome 1 and 5 region, and it is therefore plausible that they even historically have maintained larger effective population sizes.

Our analyses only favored an isolation with migration model for the region on chromosome 3, where there was migration from northern to southern populations (Figure 2). The region on chromosome 3 markedly differs from the other two regions in its geographical distribution of northern and southern haplotypes^10, 33^. While the divergent haplotypes for chromosome 1 and 5 only meet at narrow migratory divides in Europe, the contact zone for the divergent haplotypes on chromosome 3 extends from central Scandinavia eastwards to southern Siberia, which likely allows for more opportunities for gene flow. Increased gene flow in this region may also be facilitated by the apparent lack of more complex rearrangements as seen in the other two regions (Figure 2). Although not supported by simulations, we cannot rule out that there has been at least some gene flow between northern and southern populations also in the regions on chromosome 1 and 5. Double crossovers are, together with gene conversion, the main mechanism allowing for gene flow between inverted and collinear haplotypes, and are predicted to be less frequent closer to breakpoints^34^. Consistent with this prediction, we observed the highest differentiation in the vicinity of breakpoints in the regions on chromosome 1 and 5 (Figure 2).

We identified similar sequence repeats for at least some of the different breakpoints within each divergent region, which may have been directly involved in the formation of the structural changes^35^. Interestingly, arrays of the same type of tandem repeat are associated with the breakpoint regions on chromosome 1 and 3 and are not found in the corresponding intervals in the zebra finch or collared flycatcher genome. Within the northern willow warbler genome, highly similar and complete copies of this repeat (at least 50 % length and 90 % identity) are restricted to the two regions and on scaffolds predicted to be adjacent to them. It is unclear if these large tandem repeat arrays were present at the time when the structural differences were established or if they have expanded afterwards and to what extent they differ between the subspecies. The differentiated region on chromosome 5 did not show tandem repeat arrays at the breakpoints, but for two of the breakpoint intervals in the northern genome, there was a 31 kb segmental duplication containing a truncated and likely pseudogenized copy of the URB1 gene.

### Selective targets in the divergent regions

Identifying selective targets within each divergent region is challenging due to the large number of genes (N=47-197) and high linkage disequilibrium. Breakpoints themselves may be under selection if they modify the expression or disrupt the protein-coding sequence of genes^13, 36^. However, none of the breakpoint intervals overlapped with or were close to annotated functional genes (Supplementary Table 6), although we cannot rule out an effect on more long-distance regulatory elements. We identified 68 highly differentiated SNPs or short indels in 45 genes that were predicted to modify the protein-coding sequence (Supplementary Table 7). These genes are associated with a wide range of biological processes and some of them lack any functional annotation. The variant with the largest predicted impact was a frameshift deletion in the GTF3A gene located on chromosome 1, which also contains three additional highly differentiated SNPs. The frameshift variant is represented by a derived deletion in the northern willow warblers, which changes four amino acids at the end of the gene and extends the protein with three amino acids. This gene encodes a transcription factor involved in transcription of 5S rRNA genes and has in humans been associated with body mass index^37^. The highly differentiated variants in this gene could potentially be associated with physiological adaptations to the different migratory routes of the subspecies. In line with this, we also found highly differentiated nonsynonymous mutations in three fatty acid desaturase genes that are located in tandem in the region on chromosome 5. Fatty acid desaturase genes regulate unsaturation of fatty acids and have been shown to underlie dietary adaptations in humans^38, 39^. Two of these genes are associated with the gene ontology term “oxidation-reduction process”, which also includes three other genes with nonsynonymous mutations, CYB5R2 and BBOX1 on chromosome 5 and CRYL1 on chromosome 1.

The region on chromosome 3, on the other hand, shows a strong correlation with altitude and latitude in the breeding area^10, 33^ and a potential selective benefit of the northern haplotypes could be increased cold tolerance. In this case, a potential candidate gene would be LDL receptor related protein 11 (LRP11), which is annotated with the gene ontology term “response to cold”.

In order to identify additional putative functional differences, we also screened for highly differentiated structural variants. Most of these variants were located far from the closest gene and for those overlapping genes, most were located in relatively large introns. This could suggest that they are located in regions with more relaxed selection pressures and for most of them the effect on the phenotype may be minor. Two structural variants overlapped the three prime untranslated region of genes and potentially have an effect on post-transcriptional regulation: a 21 bp insertion in IFNGR1 on chromosome 3 and a 203 bp deletion in STOML3 on chromosome 1. INFGR1 plays a critical role in the immune response against intracellular pathogens^40^. The different climates associated with northern and southern haplotypes on chromosome 3 could be linked to different stress on the immune system and/or impact the abundance of pathogens. STOML3 modulates the sensitivity of mechanoreceptors^41^. Mechanoreceptors are involved in several physiological processes^42^ and the potential phenotypic effect of the structural variant is therefore difficult to predict. Overall, the functional annotation of the nonsynonymous changes and the structural variants suggest that the regions affect several different gene pathways and could potentially have widespread phenotypic effects.

## Conclusions

We have demonstrated that structural rearrangements maintain large differentiated regions despite extensive hybridization. Using a modelling approach we obtained more robust estimates of divergence times and showed that the divergent regions of the three chromosomes are of similar ages. This observation is compatible with a scenario where the inversions arose in allopatric populations that later came into secondary contact and hybridized. Finally, our improved genome and annotation has provided a set of new candidate genes for adaptations related to migration and environmental gradients.

## Methods and materials

### Samples

Nine willow warbler males were caught opportunistically with mist nets during the time of autumn migration at Krankesjön, 15 km East of Lund, Southern Sweden. The phenotypic variation (color and size) suggested that they were potentially represented by birds from both Southern and Northern Scandinavia, and thus each of the migratory phenotypes. Blood from each bird was collected through puncture of the brachial vein and was stored in two vails containing SET buffer and 70% ethanol, respectively. An aliquot of the blood was used for DNA extraction with a phenol-chloroform protocol. From the extracted DNA we genotyped the samples for two loci located on chromosome 1 and 5, respectively (NBEA and FADS2)^43, 44^, and for a bi-allelic marker within the divergent region on chromosome 3 (AFLP-ww1)^45^. Based on the genotyping results we selected two samples that were homozygous northern or homozygous southern for all three loci, respectively.

### Optical maps

DNA was extracted from blood stored in ethanol using a Bionano Plug Lysis protocol (v.30026D). The blood was first separated from the ethanol through gentle centrifugation and embedded in molten 2% agarose plugs (Bio-Rad DNA plug kit). The solidified plugs were submerged in Lysis Buffer solution (Bionano) and Proteinase K (Puregene/Qiagen, 66.8 µl per ml Buffer) for two hours at 50°C. The plugs were subsequently washed in 1x Wash buffer (Bio-Rad DNA plug kit) followed by TE buffer. In the following step, the plugs were treated with RNase (Qiagen, 20µl in 1 ml TE buffer) for 1h at 37°C, followed by another washing step using the same buffers as in the previous step. Next, the plugs were melted for 2 min at 70°C and treated with GELase (Epicenter) for 45 minutes at 43°C. The DNA was then purified from digested agarose using drop dialysis against TE buffer on a 0.1µm dialysis membrane (Millipore) for 2.5 h.

Optical maps for each of the two samples were produced using Bionano’s commercial Irys system^46^. BspQ1 was determined to be the most suitable nicking enzyme after using the software LabelDensityCalculator v.1.3.0 and Knickers v.1.5.5 to analyze a previous short-read assembly^10^. Bionano’s IrysPrep Labeling-NLRS protocol (v.30024) was used for the NLRS reaction. For this step, DNA was treated with Nt.BspQ1 (NEB) to create single-stranded nicks in a molecule-specific pattern. These were then labeled with Bionano’s labeling mix (NLRS kit), aided by Taq Polymerase (NEB), and repaired using Bionano’s repair mix (NLRS kit), in the presence of Thermopol Rxn buffer, NAD+, and Taq DNA Ligase (NEB). Finally, the DNA backbone was stained using DNA stain from Bionano’s NLRS kit. Each sample was then loaded on two IrysChips (Bionano) each, and the DNA with stained BspQ1 nicks was visualised using an Irys instrument, following Bionano’s Irys user guide (v.30047). This resulted in 200 and 182 Gb of data for the northern and southern sample, respectively.

Genome maps were assembled *de novo* using Bionano’s in house software IrysView v.2.5.1, with noise parameter set to “autonoise” and using a human arguments xml file. The genome map was then further refined by re-assembling all data, but using the first assembly version as a reference. The final assemblies were both 1.3 Gb in total size, with an average coverage of 92.3 and 96.4x, and N50 of 0.93 Mb and 0.95 Mb, respectively.

### Linked read sequencing

For the southern sample, DNA for chromium sequencing (10x Genomics) was extracted from blood stored in SET buffer using a MagAttract HMW DNAkit (Qiagen) at Scilifelab, Stockholm, Sweden. For the northern sample the extraction for bionano optical maps was used. The libraries of the northern and southern sample were each sequenced on a separate lane of a HiseqX (Illumina).

### Northern *de novo* assembly

Library preparation for long read sequencing was done on DNA previously extracted for the bionano optical map and followed Pacbio’s standard protocol for 10-20 kb libraries. No shearing was performed prior to the library construction, but the library was size selected using the BluePippin pulse field size selection system (Sage Science), with a size cut-off >25kb. The library was sequenced on eight SMRT cells on a Sequel platform (Pacbio). The sequencing yielded 63.66 Gbp of data comprised of 4,690,365 subreads with a mean length of 13,573 bp (range: 50-170,531 bp).

The pacbio reads were assembled *de novo* in HGAP4^47^ in the SMRT Link package with default settings except for specifying an expected genome size of 1.2 Gbp and setting the polishing algorithm to “Arrow”. We ran Falcon unzip^48^ on the assembly to obtain partially phased primary contigs and fully phased haplotigs. Within the software, Arrow was used to polish the assembly using reads assigned to each haplotype. The assembly was further polished with Pilon 1.22^49^ with Illumina chromium reads from the same sample.

The primary Illumina-polished scaffolds were scaffolded with linked reads using arcs v. 1.0.5^50^ and LINKS v. 1.8.6^51^ and with the optical map using bionano solve v. 3.2.2 (BioNano Genomics, CA, USA). Contigs with overlapping ends were merged in GAP5 in the staden package^52^ and contigs that could be mapped over most of their length to larger contigs and that showed limited diploid coverage were removed with the purge haplotig pipeline^53^ . We used PBJelly^54^ to close gaps and performed a second round of polishing using the chromium reads.

Repeats were annotated using Repeatmodeler version open 1.0.8^55^, repeatmasker repeatmasker version open 4.0.7^56^ and tandem repeats finder version 4.0.9^57^. We used augustus version 3.2.3^58^ to create gene models using hints provided from RNAseq data (Supplementary Table 3) and protein data from other bird species. Genes within the divergent regions were manually curated in Webapollo 2.0.4^59^. For further details on the assembly steps and annotation see the Supplementary Methods.

### Southern *de novo* assembly

We used Supernova version 2.0.0^60^ to create a *de novo* assembly from 10x chromium reads from the southern sample. The software was run with default settings and results were exported into a fasta file in a “pseudohap style” format, which contains a single record per scaffold. The assembly contained 42,450 scaffolds, with a length of 1.17 Gb and an N50 of 13.2 Mb (Supplementary Table 1). Next, we hybridized the scaffolds with bionano data from the same sample. For the hybrid assembly default settings were used with the exception of aggressive scaffolding parameters. The hybrid scaffolding made 41 cuts to the optical maps and 118 cuts to the scaffolds and generated 210 super-scaffolds and 42,291 scaffolds that could not be further scaffolded. We decided to ignore one cut that was made to a scaffold that covered the entire differentiated region on chromosome 5. The cut was made close to a large gap and was not supported by alignments of linked reads. From this assembly we removed scaffolds that were entirely comprised of Ns (due to cutting across Supernova assembly gaps with the optical map), entire duplicates of other scaffolds or that showed a substantial proportion of contaminated sequence, which resulted in a final assembly comprised of 39,597 scaffolds and with a scaffold N50 of 12.4 Mb (Supplementary Table 1).

### Whole-genome resequencing and variant calling

We used the whole-genome resequencing data from nine samples of each migratory phenotype provided in Lundberg et al.^10^ and an additional two high-coverage samples from each migratory phenotype (Supplementary Table 4). Sequencing libraries for the new samples were prepared with a TruSeq DNA PCR-Free kit (Illumina, CA, USA) with a targeted insert size of 670 bp or with a Truseq DNA nano (Illumina, CA, USA) with a targeted insert size of 350 bp. All of the new samples were sequenced on a HiseqX (Illumina, CA, USA). The raw reads were trimmed with trimmomatic 0.36^61^ with the parameters “ILLUMINACLIP:TruSeq3-PE-2.fa:2:30:10 LEADING:3 TRAILING:3 SLIDINGWINDOW:4:15 MINLEN:30”.

Quality-trimmed reads were mapped to the assembly using bwa mem with default settings except for specifying -M flag to ensure compatibility with the downstream duplicate removal steps and converted into binary alignment map (bam) files using samtools. For samples sequenced across multiple lanes, reads from each lane were mapped independently and the resulting bam files were merged with samtools. Read duplicates were removed with the markduplicates tool provided in picardtools.

From the aligned whole-genome resequencing data set we called variants with freebayes version v1.1.0 using default settings and parallelizing the analyses of separate scaffolds using gnu parallel^62^. Vcflib^63^ was used to filter the raw set of variants for sites with quality score >30 and for alternate alleles that were supported by at least one read on each strand (SAF>0 & SAR>0) and had at least one read balanced to the right and the left (RPL>0 & RPR>0). Next, we used vcftools to filter genotypes with a coverage of at least 5x and removed sites a maximum of four genotypes missing in each of the populations. The variants were also filtered for collapsed repeats by removing sites with a mean coverage of more than twice the median mean coverage (30x). We next used vcflib to decompose haplotype calls and complex alleles into indels and SNPs and removed any variants that were overlapping with annotated repeats. This gave as a final of 50 million variants of which 44 million were bi-allelic SNPs. We used vcftools to calculate F_ST_ ^64^ for each variant and for bi-allelic SNPs in non-overlapping windows of 10 kb.

Coverage for each resequenced sample was calculated in non-overlapping 10 kb windows using bedtools v2.29.2^65^ and only included properly paired reads with a mapping quality of at least 1. The raw coverage values for each sample were normalized by its median coverage across all windows.

### Structural variant calling

We used delly version 0.7.7^66^ and manta 1.0.3^67^ to call structural variants in the resequenced samples. For delly we first called variants separately in each sample, merged the variants identified across samples and re-genotyped each sample for the combined set. The raw set of variants was filtered using the germline filter included in the software with default settings except for setting no lower limit for SV size. Following filtering we obtained 67,648 deletions, 34,217 insertions, 2,120 duplications, 3,808 inversions and 34,289 breakend annotations. For manta we called variants jointly across all samples and filtered the raw data set to only include variants with a PASS flag. The filtered set of variants was comprised of 21,494 deletions, 4,168 duplications, 7,426 insertions, 2,942 inversions and 8,850 breakend annotations. Genetic differentiation (F_ST_) was calculated in vcftools and variants with F_ST_≥0.7 between homozygotes in each divergent chromosome region were extracted and checked for overlap with genes and gene features using bedtools. In the case of overlapping variants, we randomly picked one of them.

### Inversion genotypes for resequenced samples

The resequenced samples were assigned a genotype of southern and northern haplotypes for each of the divergent regions based on a multidimensional scaling (MDS)-based clustering in invclust^68^ of SNP array genotypes in Lundberg et al.^10^. To obtain genotypes of the SNPs included on the array in the resequenced samples, we mapped the SNP array probe sequences to the northern assembly using gmap and from the alignments extracted the positions of the focal SNPs. Next, we used freebayes to genotype the resequenced samples for these positions and plink version 1.9^69^ to combine the genotypes with the genotypes from the SNP array. In the genotyping step we also included mapped 10x chromium libraries for the northern and southern reference samples (see below) to be able to confirm that these samples were homozygous northern and southern for all of the divergent regions, respectively. From the combined dataset we extracted genotypes for SNPs located in each of the divergent regions and used invclust to assign each sample a genotype of inverted and non-inverted haplotypes. The inverted and non-inverted haplotypes were recoded as southern or northern haplotypes based on their frequency in each subspecies.

### Breakpoint analyses

We used SatsumaSynteny to align the genomes of the southern and northern willow warblers, and the northern willow warbler genome to the genomes of the zebra finch and collared flycatcher FicAlb 1.5^70^.

To provide further evidence of breakpoints, we mapped the 10x chromium reads of each sample to both the northern and the southern assembly and called structural variants using the longranger wgs pipeline. For the southern genome we selected the 500 largest scaffolds to make it compatible with the software. We also checked for differences in linked read molecule coverage between the samples. For this purpose, the raw reads of each sample were first processed with longranger basic for quality trimming and barcode processing. The trimmed reads were mapped to the assemblies using bwa mem using a -C flag to extract the barcode information of each read and alignments converted into bam files using samtools. To estimate coverage of barcodes we first used the tigmint-molecule script from tigmint 1.1.2^71^ to obtain positional information of barcodes (molecules) in each divergent region. The software was run with default settings except for only using reads with a mapping quality of at least 1 and only to report molecules that were estimated to be at least 10 kb. We next used bedtools to count the number of overlapping molecules in 1 kb windows.

We explored differences between optical maps by using the runSV.py script in bionano solve with the southern optical map as a query and the northern assembly as target and the reciprocal analysis with the northern optical map as a query and the southern assembly as a target. We also used the bionano solve hybrid assembly pipeline to visualize differences between the optical maps and the genome assemblies at breakpoint regions.

### Functional annotation of differences

We used bedtools to quantify the distance between breakpoint intervals and annotated genes. To provide a functional annotation of the SNPs and short indels, we selected variants that showed a F_ST_≥0.7 between southern and northern homozygotes for each of the region and used as input to snpeff together with the annotation and reference genome^72^. Snpsift^73^ to select variants that were predicted to have a moderate to high effect on genes. Gene ontology terms for the genes were extracted from the chicken orthologues (ensembl) or through domain searches of the proteins with interproscan.

### Age estimation of divergent regions

In order to estimate the timing of the inversion events we used high-coverage resequencing data from two *trochilus*, two *acredula* and, as an outgroup, one dusky warbler *Phylloscopus fuscatus* (Supplementary Table 4). The willow warbler samples were chosen so that they were either homozygous southern (*trochilus*) or northern (*acredula*) for all of three divergent regions. The dusky warbler library was prepared using a TruSeq Nano DNA library prep kit for Neoprep (Illumina) according to the instructions of the manufacturer and sequenced on a Hiseq X. Quality-trimming of the raw reads and mapping of the trimmed reads to the northern reference genome followed the same approach as used for the willow warbler resequencing samples (see above).

Variants were called using freebayes and the raw set of variants were filtered using gIMble’s preprocess module (v0.6.0). Sample-specific callable sites were identified using gIMble preprocess and were defined as those with a minimum coverage of 8x and a maximum of 0.75 standard deviations above the mean coverage. Genic and repetitive regions of the genome were removed from the callable sites in order to limit downstream analyses to intergenic regions.

Summary statistics of genetic variation (π and d_xy_) within the divergent regions were calculated using gIMble. Following this, net divergence (d_a_) between northern and southern samples was calculated as d_north-south_ - (π_north_ + π_south_)/2. To convert the net divergence into years we used the germline mutation rate (4.6×10^-9^) estimated in the collared flycatcher^23^. Relative node depth (RND) using the dusky warbler (DW) as an outgroup was calculated as d_north-south_/(d_DW-north_+d_DW-south_)/2. For each divergent region a blockwise site frequency spectrum (bSFS) was generated with gIMble using blocks of 64 bp in length. This length refers to the number of callable sites within a block, while the physical length of blocks was allowed to vary due to missing data but was limited to 128 bp. Downstream analyses that relied on a bSFS used a kmax of 2, meaning that only marginal probabilities were calculated for mutation counts >2. The composite likelihood (CL) of a model, given the bSFS of one of the divergent regions, was optimized using the Nelder-Mead algorithm with the maximum number of iterations set to 1000. Within the software we evaluated three different population models. The first model was a strict isolation model (SI), with parameters ancestral effective population size, effective population sizes for southern and northern willow warblers and divergence time. The second model was an isolation with migration model (IM_1_) that also included a migration rate from northern to southern samples and the third model (IM_2_) instead had a migration rate from southern to northern willow warblers.

Simulations were carried out by msprime 0.7.4^74^ through gIMble. The recombination rates used for these simulations were chromosome-specific estimates from a high-density recombination map of the collared flycatcher^70^ and were 2.04, 1.95, and 2.63cM/Mb for chromosomes 1, 3, and 5 respectively. A total of 100 replicates were simulated for the optimized SI parameters of each region. These simulated bSFSs were then optimized under both an SI model as well as the best fitting IM model for that region. The improvement in CL between these models was used as a null distribution for testing whether improvements in CL observed for the real data were greater than expected given a history of no migration. For each parameter we calculated 95 % CI as MCL estimate ± 1.96 * standard deviation of simulations (Supplementary Table 9). As a result, our estimates of uncertainty are affected by the recombination rates that we assumed for simulations. We also used the results of simulations to quantify the potential bias in MCL estimates due to intra-block recombination (Supplementary Table 9). However, we do not attempt to correct for this bias as it is relatively small (e.g. the MCL divergence times are estimated to be biased upwards by 8, 12 and 19%) and our estimation of the bias itself is largely dependent on the recombination rates we assumed.

MSMC2^26^ was used to explore genome-wide changes in Ne through time. As input to the software we used the callable intergenic bed file and filtered vcf file mentioned above, with the addition of further filtering the bed file to only include autosomal scaffolds ≥ 500kb and excluding the divergent regions. The input files for MSMC2, i.e. an unphased set of heterozygous sites for each sample, were generated using the generate_multihetsep.py script from msmc-tools. MSMC2 was run with a starting ρ/μ of 1 for 30 expectation-maximum iterations. For both the demographic modelling and MSMC2 we used the collared flycatcher germline mutation rate^23^ and a generation time of 1.7 years^8^ to convert divergence times into years.

## Supporting information

Supplementary methods and results

## Data accessibility

Raw sequence data, optical maps and *de novo* assemblies are available at NCBI under bioproject PRJNA550489. Annotation files are available at https://github.com/maxlundberg/warbler_inversions.

## Code availability

Workflows and scripts are available at https://github.com/maxlundberg/warbler_inversions.

## Author contributions

ML and SB conceived the study. ML performed assembly curation and further scaffolding of the genome, annotation, supervised other bioinformatics analyses and wrote the manuscript with input from the other authors. AM performed the dating analyses. AP performed extraction of the northern sample used for bionano optical maps and pacbio sequencing, performed the pacbio *de novo* genome assembly and early analyses of the bionano data.

## Acknowledgements

We would like to thank Konrad Lohse and Lars Råberg for comments on the manuscript, Sarah Kingan for discussions on pacbio assemblies in highly heterozygous species, as well as Gertjan Bisschop and Dominik R. Laetsch for their advice on the gIMble analysis. The study was supported by grants from the Royal Physiograhic Society of Lund to ML and by grants from the Swedish Research Council (2017-03937) and Crafoord Foundation to SB. The authors would like to acknowledge support from the National Genomics Infrastructure (NGI) in Uppsala (Uppsala Genome Center/SNP&SEQ Technology Platform) and in Stockholm. Work performed at NGI has been funded by RFI/VR and the Science for Life Laboratory, Sweden. Computations and data handling were enabled by resources provided by the Swedish National Infrastructure for Computing (SNIC) at UPPMAX, partially funded by the Swedish Research Council through grant agreement no. 2018-05973.

## Notes

### Competing Interest Statement

The authors have declared no competing interest.

