## Supplementary methods and results for "Inversions maintain differences between migratory phenotypes of a songbird"

##### Assembly of a northern willow warbler genome

The pacbio reads were assembled *de novo* in HGAP4[1] in the SMRT Link package with default settings except for specifying an expected genome size of 1.2 Gbp and setting “Arrow” as the polishing algorithm. We ran Falcon unzip[2] on the assembly to obtain partially phased primary contigs and fully phased haplotigs. Within the software, Arrow was used to polish the assembly using reads assigned to each haplotype.

We evaluated two unzipped assemblies based on 30x or 40x coverage of seed reads in the preassembly step in HGAP4. A lower coverage threshold will lead to longer reads in the initial assembly step, which may increase the contiguity of the assembly, but will on the other hand limit the number of reads that can be used in the phasing and polishing step. Although the unzipped assemblies were very similar, the 40x version was chosen for downstream analyses as it was slightly more contiguous and contained a higher number of single-copy bird orthologues as determined by BUSCO version 3.0.2[3].

The assembly was further polished using Pilon 1.22[4] with Illumina chromium reads (2x150 bp) from the same sample. The Illumina reads were mapped to the genome using bwa version 0.7.17-r1188[5] and duplicated reads were marked using picardtools 2.10.3 (http://broadinstitute.github.io/picard). Pilon was run by only correcting indels and in total the software made 1,043,827 insertions and 275,457 deletions, respectively, of which the vast majority (94 %) were single basepair changes. The Illumina polishing had a pronounced effect on the number of single-copy bird orthologues that could be detected in the primary contigs (Supplementary Table 1).

For further assembly steps we extracted the Illumina-polished primary pacbio contigs (N=2,737, N50 of 2.1 Mb and a length of 1.29 Gb). These contigs showed an unexpectedly high level of duplicated single-copy orthologues (7.4%), which suggested partial or complete overlap between some contigs. As a first step to reduce the redundancy and increase the contiguity of the assembly we hybridized the primary contigs to the bionano optical map of the same sample using bionano solve version 3.2.2 (BioNano Genomics, CA, USA) with default settings except for specifying aggressive scaffolding parameters. The hybrid scaffolding resulted in 19 cuts to the bionano maps and 259 cuts to the pacbio contigs and created 363 super-scaffolds. Most of the gaps between the contigs in the super-scaffolds were estimated to be negative. However, in the hybrid assembly, sequences on either side of these gaps were not collapsed and thus formed false segmental duplications. To remedy this problem we extracted 304 sets of overlapping contigs ("supercontigs") and used GAP5 in the staden package[6] to find potential joins between the contig ends. Using this approach we merged contigs at 558 (87%) of the putative overlaps. The mean alignment length in the overlaps was 111 kb (range: 0.259 – 661 kb) with a mean sequence divergence of 3.28 % (range: 0.31-15.55 %). The highest divergence was caused by the presence of large indels. By trimming off one or both ends of the contigs at the gaps (mean 23 kb, range: 0.6-60 kb), we were able to close 23 further gaps. For the remainder of gaps, GAP5 failed to find potential joins between contigs or the ends supposed to be joined were considered to have too high divergence. The new assembly including supercontigs consisted of 2,401 contigs with an N50 of 6.5 Mb and had a considerably lower amount of duplicated single-copy genes (4.6 % vs 7.4%).

To further reduce the redundancy we used the purge haplotig pipeline[7] to remove contigs that could be mapped over most of their length to larger contigs and that showed limited diploid coverage. We first estimated coverage by mapping the pacbio subreads used for the *de novo* assembly with minimap2 version2.13-r860[8] using default settings for pacbio reads (-x map-pb). To minimize the loss of repetitive sequences that could be separated and scaffolded by the bionano optical map, we used the first bionano hybrid assembly (363 superscaffolds and 1500 cut and unscaffolded contigs) as a reference for mapping. From the mapped data we detected a clear haploid and diploid peak and set a threshold of diploid coverage above 34x and below 85x. Any scaffold where less than 80 % of its positions had diploid coverage was considered a putative haplotig and was mapped to other scaffolds using minimap2 within the software. We removed 1209 scaffolds (mean size: 107,655 bp, range: 598-495,788 bp) with a coverage to the best hit of at least 70 % (mean: 97.4 %). Using this approach we specifically excluded contigs that could not be incorporated in superscaffolds. However, we also removed three contigs that each entirely made up short superscaffolds that could be uniquely assigned to larger superscaffolds and that had a high degree of haploid coverage. At this stage we also removed five additional contigs shorter than 1000 bp that were the result of cutting the assembly with the bionano optical map. This led to an assembly with 1187 contigs, a length of 1.1 Gbp and a N50 of 7.9 Mb. The filtered assembly showed a large reduction in single-copy orthologue bird genes (1.3 vs 4.6%).

To provide an intermediate level of scaffolding to the optical map, we mapped the 10x chromium reads of the same sample to the assembly using bwa and used arcs version 1.0.5[9] and LINKS version 1.8.6[10] for scaffolding. Arcs was run with default settings except for enabling gap size estimation (--dist_est) and LINKS was run by setting the number of supporting links to at least 5 (-l=5) and the maximum link ratio between the two best contig pairs to 0.3 (-a=0.3). The scaffolding resulted in 739 scaffolds with a N50 of 16.4 Mb and a length 1.12 Gb.

As a final scaffolding step we hybridized the 10x chromium-pacbio scaffolds to the bionano optical map using the same settings as before. The hybrid scaffolding made 23 cuts to the optical map, 122 cuts to the scaffolds and resulted in 497 scaffolds with an N50 of 16.8 Mb. Two contigs representing the divergent region on chromosome 1 had been scaffolded together by arcs but were separated and not re-scaffolded with other sequences in the bionano hybrid assembly. Since the mismatched end of the optical map was short, located at a large gap and the gene order is the same as seen in other bird genomes, we decided to keep the scaffold generated by arcs.

For this round of hybrid scaffolding, there were 52 gaps that were estimated to be negative (i.e. some overlap between scaffolds). Using the same approach as when creating supercontigs, we were able to close 10 of these gaps. We additionally closed gaps using PBJelly[11] with default settings except for specifying --spanOnly --capturedOnly". The software filled 97 gaps, extended one end of 12 gaps, extended both ends of 18 gaps and overfilled 28 gaps.

We further checked for potential misjoins between scaffolds that originate from different chromosomes. To this end, we used SatsumaSynteny[12] to produce whole-genome alignments between the assembly and the genomes of chicken (version GRCg6a) and zebra finch (version taeGut3.2.4), both downloaded from Ensembl (www.ensembl.org). Using this approach we detected a scaffold that showed good alignments to both chromosome 10 and 23 in both of the other species. We considered this join unlikely and decided to split the scaffold.

Next, we performed a second round of polishing with the 10x chromium Illumina data from the same sample. For this round, since we had fewer than 500 scaffolds, we used the longranger 2.1.14 align pipeline[13] to map reads in a barcode-aware way. Pilon was then run with the same settings as before and resulted in the correction of 417,032 indels, of which 78.7% were single-basepair changes. The second round of polishing considerably increased the number of single-copy bird orthologues that could be identified in the assembly (Supplementary Table 1).

The mitochondrial genome was not found in the original pacbio genome assembly. We obtained this genome by adding the complete mitochondrial sequence from a previous short-read assembly[14]. We then used bwa to map the 10x chromium reads from the northern sample to the assembly and extracted alignments on the mitochondrial sequence. Next, freebayes was used with a haploid setting to detect differences present in the aligned reads. The raw variant file was filtered with vcftools for sites with a quality less than 30 and for two intervals with excessive read coverage. The filtered variant file contained 11 substitutions and three indels, and was used with bcftools version 1.10[15] to create a new mitochondrial reference.

For the extraction and removal of sequences in the different assembly steps we used kentUtils (https://github.com/ucscGenomeBrowser/kent). Summary statistics for each assembly (e.g. N50) were calculated using the assemblathon_stats.pl script[16].

**Repeat annotation**

We used Repeatmodeler version open 1.0.8[17] for *de novo* identification of repeats in the assembly. The repeats detected by repeatmodeler were combined with 1,023 bird-specific repeats into to a custom library. We used repeatmasker version open 4.0.7[18] with the custom library and by using a more sensitive search (-s flag) to annotate repeats in the genome. Next, bedtools together with the annotated repeats was used to create a softmasked version of the assembly from the northern sample, which was used in the gene annotation step. The same repeat library was also used to annotate repeats in the *de novo* assembly of the southern sample. Intervals with tandem repeats in divergent regions were also analyzed with tandem repeats finder version 4.0.9[19] using default settings except for specifying a maximum period size of 2000 bp.

Duplicated intervals within the divergent scaffolds were identified with Minimap2 and subsequently aligned with EMBOSS Stretcher (https://www.ebi.ac.uk/Tools/psa/emboss_stretcher/).

**RNA sequencing**

We used total RNA extracted from whole brain from six samples used in an earlier study quantifying differential expression in migratory and breeding willow warblers[20] (Supplementary Table 3). The quality of the RNA was checked with a Bioanalyzer version 2100 (Agilent, CA, USA). All of the extractions had a RNA Integrity Number (RIN) of at least > 7.10. RNA libraries for sequencing were prepared using a TruSeq Stranded mRNA Sample prep kit with 96 dual indexes (Illumina, CA, USA) according to the instructions of the manufacturer with the exception of automating the protocols using an Agilent NGS workstation (Agilent, CA, USA) using purification steps as described in Lundin et al.[21] and Borgström et al.[22]. The raw RNA data was trimmed using cutadapt version 1.8[23] within Trim Galore version 0.4.0 (https://github.com/FelixKrueger/TrimGalore) with default settings.

We used Stringtie version 1.3.3[24] to create transcripts from the RNAseq data. These transcripts were not used directly in the generation of gene models, but used in the manual curation step as potential alternative transcripts. For the software we first mapped the reads with Hisat2 version 2.1.0[25] using settings default settings for stranded sequence libraries and downstream transcript analyses.

**Generation of gene models**

We used augustus version 3.2.3[26] to create gene models using hints provided from RNAseq data and protein data from other bird species. For the RNAseq data we mapped the trimmed reads to the assembly using gsnap version 2018-07-04[27]. Accessory scripts in the augustus package were used to filter the alignments for paired and uniquely mapped reads and for extracting intron hints. We additionally generated coverage wig files for each strand from the filtered alignment file using the software stranded-coverage (https://github.com/pmenzel/stranded-coverage) and used these as input for the augustus wig2hints.pl to generate exonpart hints.

For homology evidence we downloaded a set of bird proteins from Ensembl (www.ensembl.org; Ensembl release 95). This data set included 28,345 proteins (16,779 genes) from chicken, 18,204 proteins (17,488 genes) from zebra finch and 15,983 proteins (15,303 genes) from collared flycatcher. We also downloaded an additional dataset from Uniprot (www.uniprot.org) that consisted of 3,169 manually reviewed bird proteins and 177 and 11,890 bird proteins that were not manually reviewed but supported by protein or transcript data, respectively. The protein data was mapped to the genome using exonerate version 2.4.0 [28]. We used the script align2hints.pl from braker 2.1.1 to generate CDSpart, intron, start and stop hints from the data.

Augustus was run with species-specific parameters (see training augustus below) and with default settings except for specifying "softmasking=true", "--alternatives-from-evidence=true", "--UTR=on", "--gff3=on" and "--allow_hinted_splicesites=atac". In the extrinsic configuration file we changed the malus for introns from 0.34 to 0.001, which increases the penalty for predicted introns that are not supported by the extrinsic data (RNAseq and protein hints). The prediction resulted in 24,140 genes and 30,316 transcripts.

**Filtering of raw models and functional annotations**

The augustus-derived gene models were annotated based on overlap with synteny-transferred chicken genes. For this purpose we used satsumasynteny with default settings to obtain whole-genome alignments between our assembly and the chicken genome. Based on the alignment, we used kraken [29] to transfer the chicken genome annotations. We then extracted the CDS from the augustus gene models and the kraken genes and used bedtools intersect to quantify the overlap. The gene models were also searched against the longest translation of each of the 16,779 chicken genes used as evidence for the gene prediction step and against 85,546 swissprot vertebrate proteins using blastp 2.5.0+[30] with an E value threshold of 1e-5. Gene models that were not annotated through synteny were assigned a gene name based on the blast results. Protein domains in the gene models were annotated with interproscan v 5.30-69.0[31]. To reduce the number of false positive predictions we removed 5,191 genes that were not supported by synteny to chicken genes, showed no significant similarity to vertebrate proteins or did not contain any annotated protein domains. To obtain a more complete annotation of the mitochondrion, we used the MITOS Webserver[32].

We used Webapollo 2.0.4 to manually curate gene models in the previously identified divergent chromosome regions and in other regions where differences were present. In the curation step we specifically validated the support for the coding sequence and the UTR and also removed genes that were likely to be pseudogenes based on a truncated coding sequence compared to homologous genes in other vertebrates, had no support from synteny in other bird species and/or that were located in repeat-rich regions. Several of the genes in the divergent region had through the automated annotation only received an ensembl gene identifier (e.g.

ENSGALG00000012194) but no associated gene name. In this case, we obtained a gene name based on the NCBI annotation of the same genome (https://www.ncbi.nlm.nih.gov/genome/gdv/browser/genome/?id=GCF_000002315.6) or through the best hits to swissprot vertebrate proteins.

**Training augustus**

We used a previous repeat-masked short-read assembly[14] and the trimmed RNAseq data used in this study to obtain species-specific parameters for augustus. The RNAseq data was assembled into transcripts using Trinity version 2.0.2[33] to create a *de novo* and a genome-guided assembly that together were comprised of 1,929,396 transcripts. The genome-guided transcript assembly was based on RNAseq mapped to the genome using GSNAP with default settings. We used PASA version 2.0.2[34] to create high-quality transcripts. The pasa transcripts were imported into Webapollo. To assess the completeness of the transcripts we compared them to synteny-transferred models from the chicken genome using kraken. We selected 1249 transcripts that appeared complete, were not overlapping with other genes and showed less than 80 % amino acid similarity to another gene in the training set. From this set we excluded 21 genes that were giving initial training errors, which gave us training set of 1228 genes. This gene set was randomly split into 1028 training genes and 200 genes used for testing. For training we used the optimize_augustus.pl script with default settings except for the flag –UTR=on.

### Supplementary results

**Chromosome 1 region**

To get further support for the structural differences observed with the alignments between the northern and southern scaffolds in the chromosome 1, we used the longranger wgs pipeline to call structural variants in linked reads aligned to either assembly. The software did, however, not report any high-confidence structural variants that overlapped the breakpoint regions. As a complementary method to detect breakpoints, we estimated the coverage of linked read molecules for each sample in 1 kb windows. As predicted, we found a dramatic drop in barcode coverage unique to the northern sample at the breakpoint region in southern scaffold (Supplementary Figure 3; Scaffold11 mid part). We also explored whether the optical map data in itself supports the structural differences between the subspecies. Using the northern assembly as a target, a translocation from the start to end of the scaffold was detected in the southern sample, which is consistent with the arrangement seen in the southern assembly.

There were also several cases where parts of the divergent scaffold or its predicted adjacent scaffolds in the southern assembly could not be aligned to the corresponding regions in the northern assembly. In the southern assembly, the two inversions are connected with a 9.8 kb sequence that is absent in the northern assembly but that would be expected in the 10 kb gap between the end of the differentiated scaffold (Scaffold156) and the scaffold predicted to be downstream of it (Scaffold143) based on the gene order in other bird species (Supplementary Figure 2). Alignments of linked reads of the northern sample to this part of the southern scaffold suggests that this part is also present in the same location in the northern species (Supplementary Figure 3; Scaffold11 mid part, between dashed lines). In addition, the last 9 kb of the divergent scaffold in the southern assembly did not align to the differentiated region in the northern genome (Supplementary Figure 2). The best match of this interval in the northern assembly is to a scaffold that is placed towards the end of chromosome 1 in zebra finch and flycatcher, although, almost equally good matches are found in several other places in the genome. When aligning linked reads to this scaffold in the southern genome, the northern sample shows a much more pronounced drop in barcode coverage than the southern sample (Supplementary Figure 3; Scaffold11 end), which could suggest the presence of another breakpoint. Finally, in the scaffold predicted to be upstream of the divergent scaffold in the southern genome (Scaffold74), the start extends 2 kb further than in the corresponding scaffold (Scaffold374) in the northern assembly (Supplementary Figure 2). The end of this interval contains 250 bp of the same type of the tandem repeat that is associated with the differentiated region, preceded by a 563 bp interval containing a partial LINE/CR1 repeat (333 bp) showing 97 % identity to the start of the divergent region in the northern assembly. Although not included in the northern assembly, alignments of linked reads to the southern assembly suggest that it is also found in the same location and orientation in the northern sample.

**Chromosome 5 region**

The breakpoint between the inverted and collinear intervals, occurring at ~ 4.0 Mb in the northern scaffold (Scaffold68), was detected by the longranger wgs pipeline in the linked read data and in the optical map data using the structural variant module in bionano solve.

Alignments of linked reads confirmed the presence of a breakpoint at the start of the divergent region (Supplementary Figure 3; Scaffold68 400 kb region). In this case, the southern sample shows a dramatic drop in barcode coverage, which is not observed in the northern sample. This breakpoint is also supported by the optical map of the southern sample, which shows a mismatch in nicking sites in this region compared to the northern assembly (Supplementary Figure 4).

The optical map data also demonstrated the presence of another breakpoint occurring just downstream of the gap in the northern scaffold (~4.1 Mb), beyond which the optical map of the southern sample continues for several 100 kbs with no matches to the optical map of the northern assembly (Supplementary Figure 4). This breakpoint was also suggested by the alignments of linked reads (Supplementary Figure 3; Scaffold68 4 Mb region). The alignments of the reads of the southern sample extend beyond the alignment of the corresponding scaffold in the southern *de novo* assembly, but the coverage drops to zero well before the northern sample.

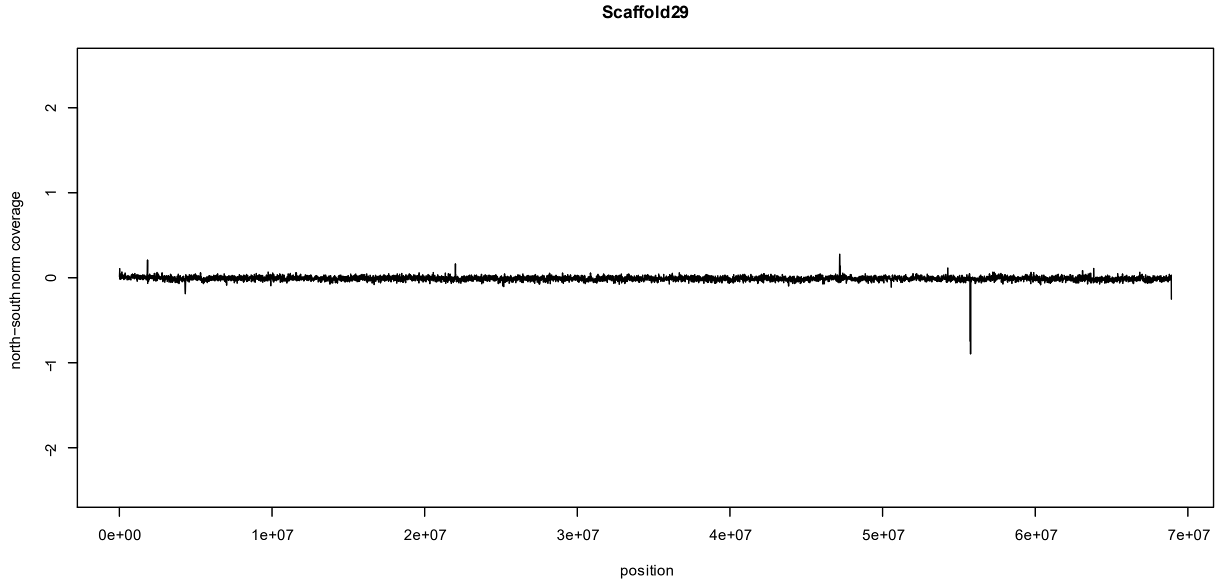


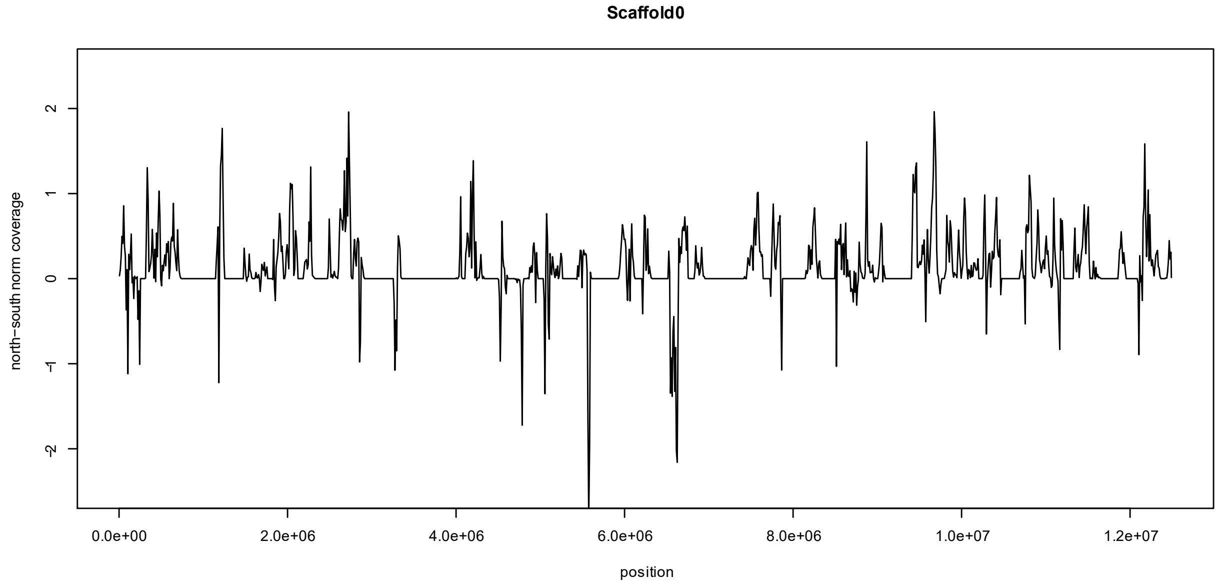


**Supplementary Figure 1**. Difference in mean normalized coverage (norm $cov_{north}$-norm $cov_{south}$) between 11 northern and 11 southern resequenced willow warblers for two scaffolds in the northern assembly. Scaffold29 contains the divergent region on chromosome 3. Scaffold0 is a repeat-rich scaffold that contains highly differentiated variants between the willow warbler subspecies, but it has not been assigned to a specific chromosome. Coverage has been calculated in non-overlapping windows of 10 kb and only for properly paired reads with a minimum mapping quality of 1.

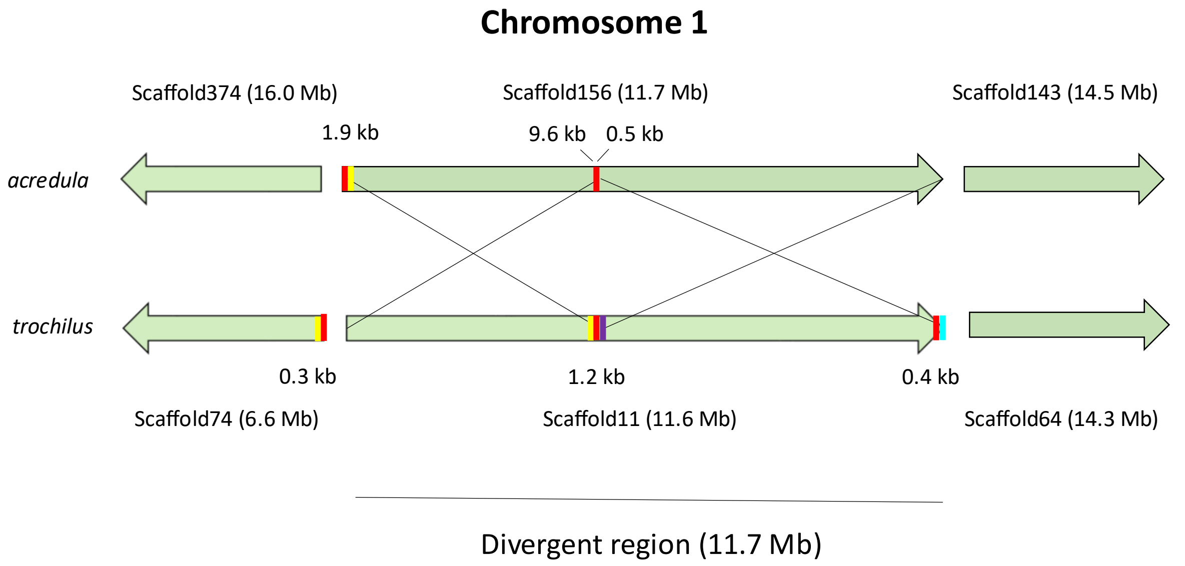


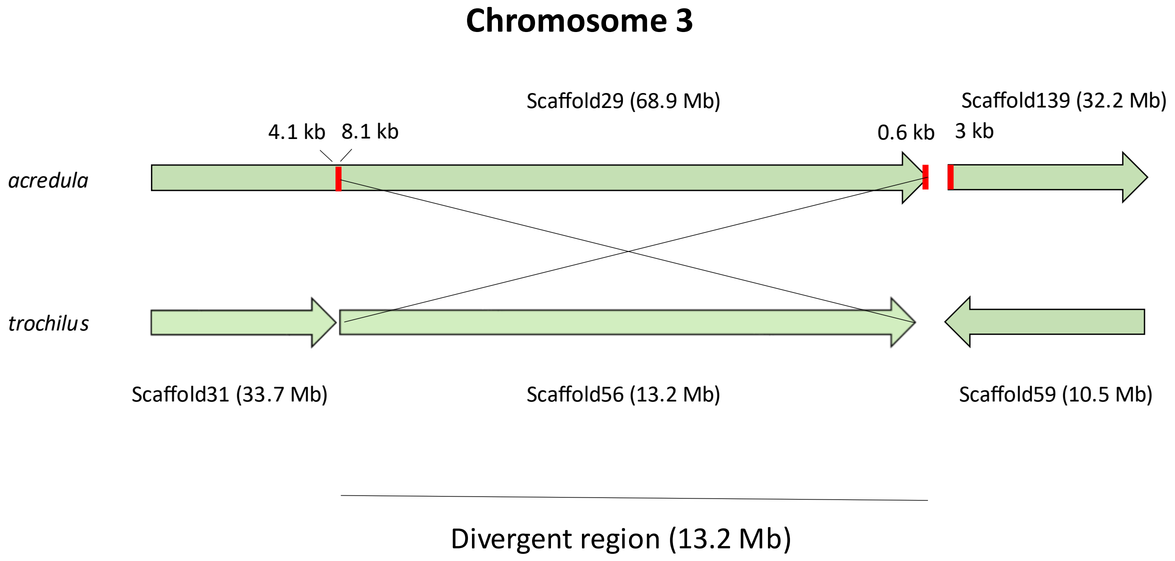


**Supplementary Figure 2**. Divergent scaffolds and predicted adjacent scaffolds for the region in chromosome 1 and 3 in the northern (*acredula*) and southern (*trochilus*) assembly, respectively. Arrays of a 413 bp tandem repeat are marked in red, with sizes presented in kb. Outside of the divergent regions, only arrays at the ends of the scaffolds have been marked. Yellow denotes a

563 bp interval downstream of the divergent region in the southern genome that shows 97 % identity to the start of the start of the divergent region in the northern genome. The same duplicated sequence is also found in the southern genome (also 97 % identity) but only a 455 bp alignment was possible due to the presence of a gap. Purple denotes a 9.8 kb interval that has not been assembled in the northern genome but that is expected at the end of the divergent region. Cyan represents a 9.0 kb interval that is present at the very end of the divergent scaffold in the southern genome but that is lacking in the divergent scaffold in the northern genome.

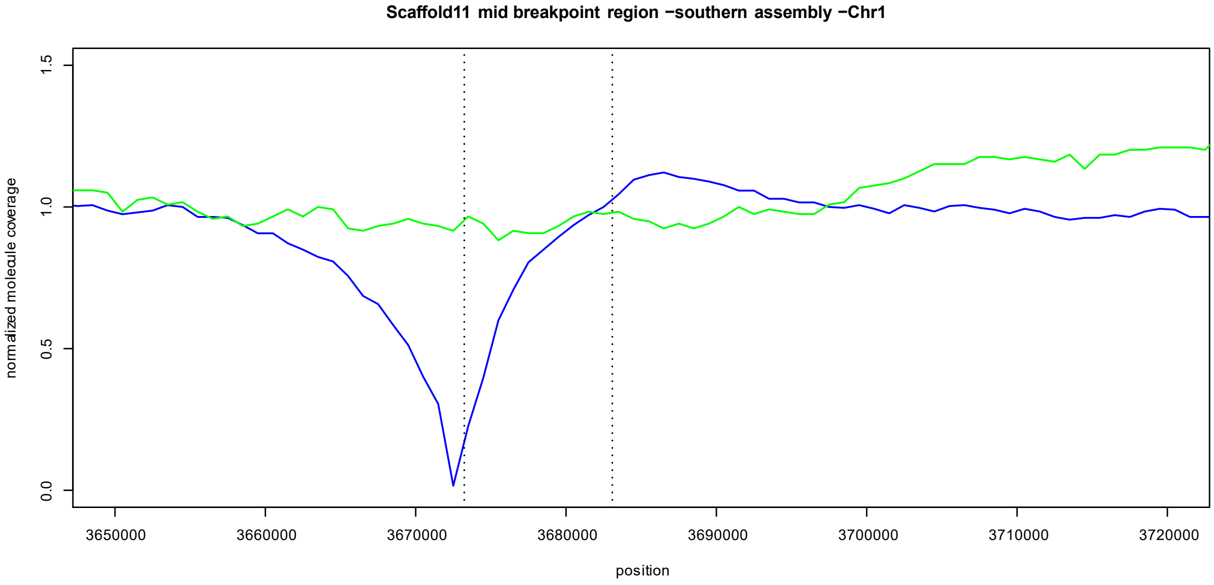
Scaffold11 mid breakpoint region −southern assembly −Chr1
normalized molecule coverage
0.0
0.5
1.0
1.5
3650000
3660000
3670000
3680000
3690000
3700000
3710000
3720000
position

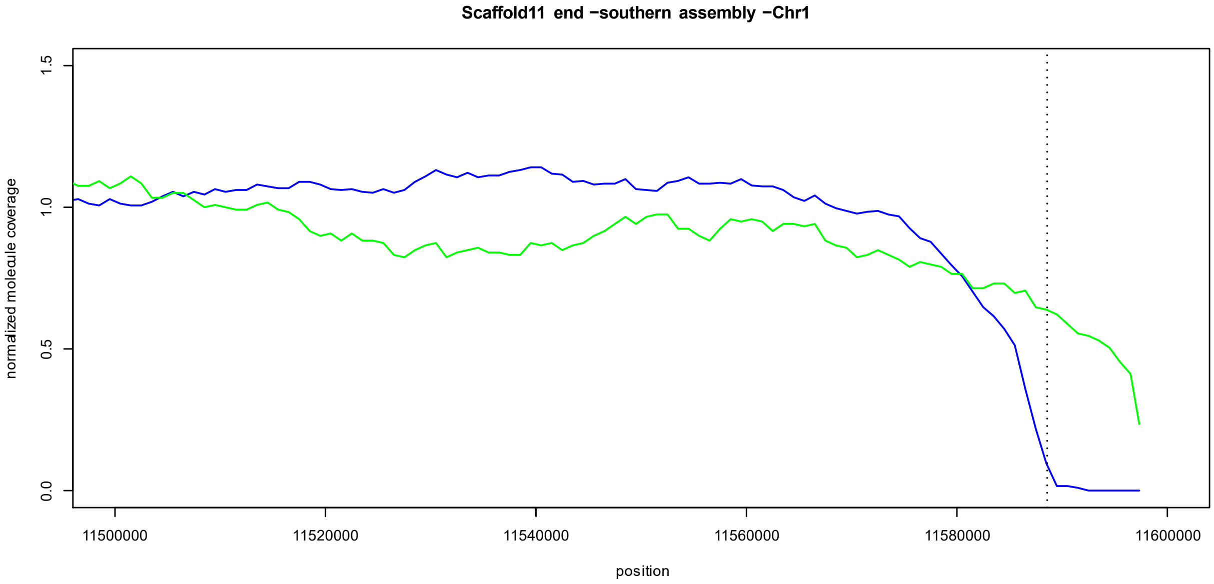
Scaffold11 end −southern assembly −Chr1
normalized molecule coverage
0.0
0.5
1.0
1.5
11500000
11520000
11540000
11560000
11580000
11600000
position

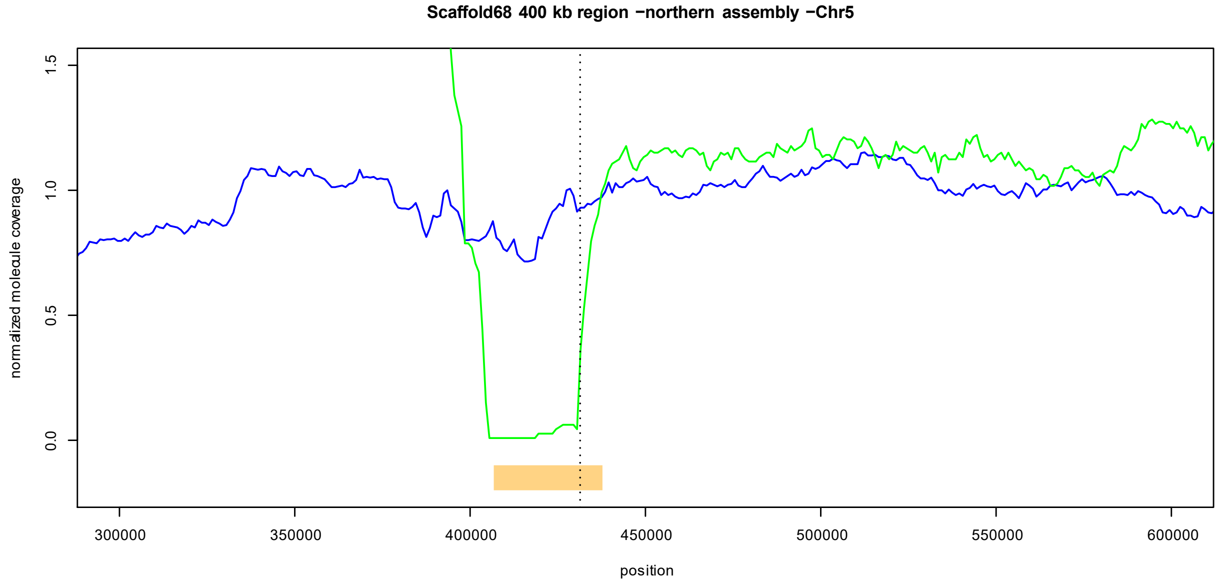


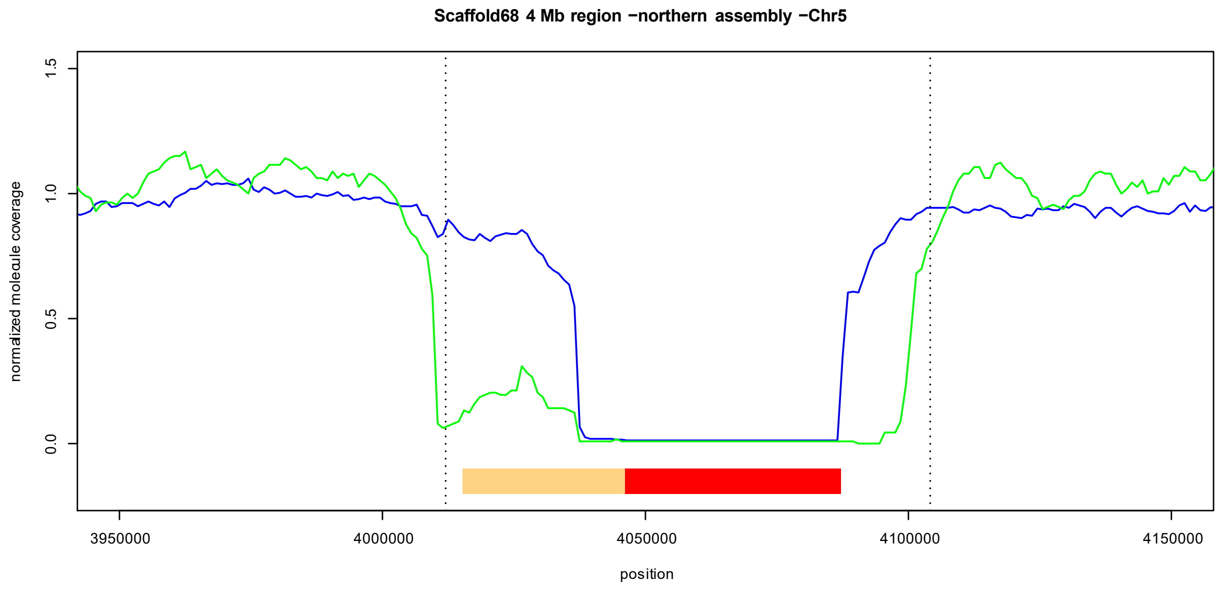


**Supplementary Figure 3**. Normalized 10x chromium molecule coverage for the southern (green) and northern (blue) samples in breakpoint regions. Normalized molecule coverage refers to the number of 10x chromium barcodes (molecules) present in non-overlapping 1 kb windows, which has been normalized by the median 1 kb molecule coverage for the sample in each of the divergent regions. Only barcodes associated with molecules that are at least 10 kb have been included. Dashed lines refer to the endpoints of the alignments of scaffolds from the assembly of the other subspecies. For Scaffold68, 31 kb duplicated intervals and a 41 kb gap are highlighted in orange and red, respectively.

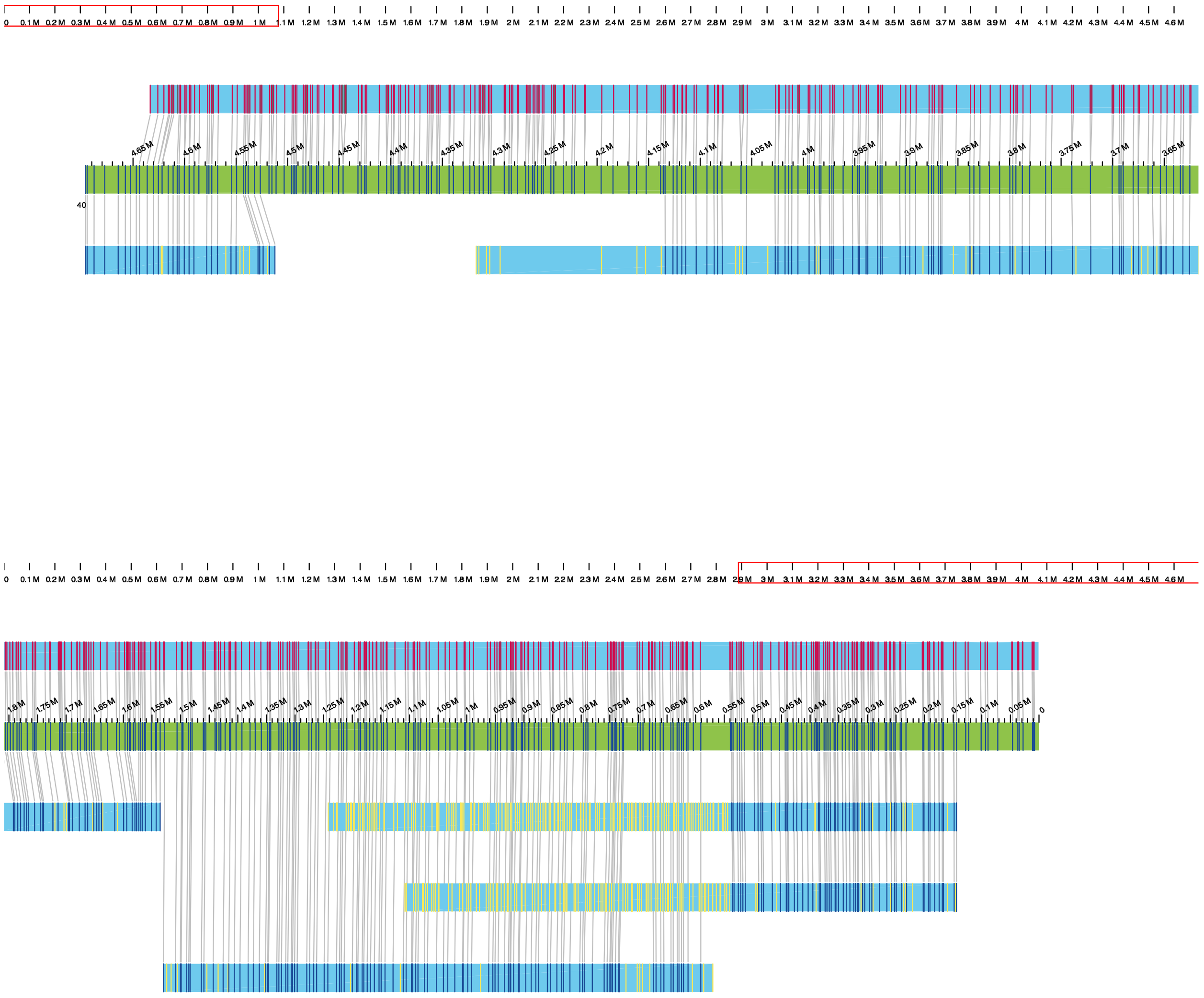

**Supplementary Figure 4**. Hybrid assemblies between the southern optical map and the northern assembly. The first and second panels show the start and the end of the divergent region on chromosome 5, respectively. In each panel, the upper blue segment represents the optical map of the northern assembly (blue), the green segment the hybrid assembly between the northern assembly and the southern optical map and the blue segments at the bottom, the optical map (contigs) of the southern sample. Matching nicking sites between the southern optical map and the northern genome are connected with vertical lines and highlighted with black bars in each contig. Yellow bars within contigs represent unmatched nicking sites. To visualize the mismatched ends of the southern optical map, it was hybridized without allowing it to guide cuts in the northern assembly.

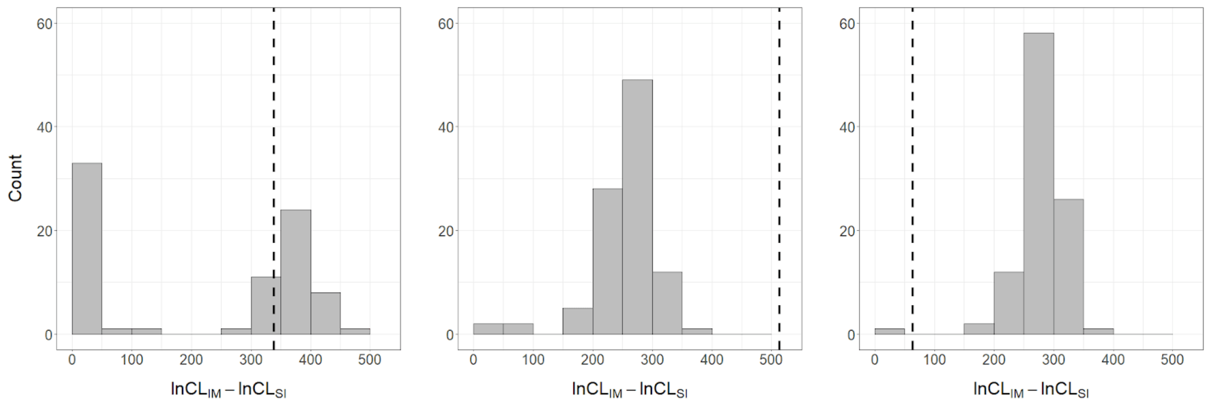


**Supplementary Figure 5.** Histograms of improvement in composite likelihood (CL) when optimizing 100 simulated strict isolation (SI) datasets under an isolation with migration (IM) model rather than SI. Plots, left to right, are for the chromosome 1, 3, and 5 regions respectively. The dashed lines represent the improvement in the real data, showing that only the chromosome 3 region has an improvement in CL which is greater than expected when there is no migration.

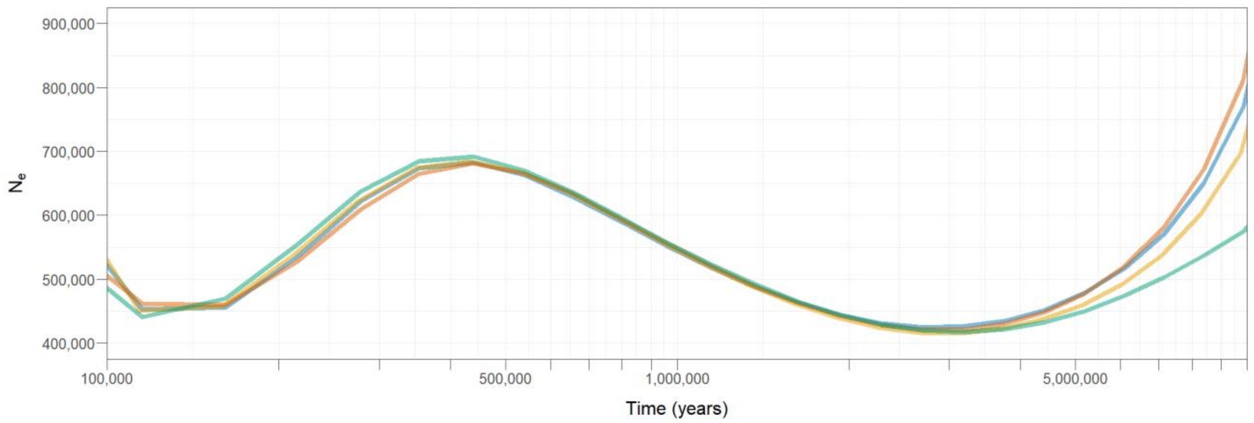


**Supplementary Figure 6**. Estimates of genome-wide $N_e$ over time inferred from MSMC2 for four high-coverage willow warbler samples. For the analysis intervals overlapping the divergent regions as well as scaffold shorter than 500 kb have been excluded. Time, between 100KYA and 10MYA, is plotted on a log10 axis.
